# Trichinella spiralis secretes stage-specific extracellular vesicles for modulating host microenvironment for parasitism

**DOI:** 10.1101/2023.08.02.551739

**Authors:** Sukhonthip Khueangchiangkhwang, Zhiliang Wu, Ritsuko Hosoda-Yabe, Tomio Yabe, Toru Okamoto, Chihiro Takada, Hideshi Okada, Shinji Ogura, Yoichi Maekawa

## Abstract

Extracellular vesicles (EVs) regulate cell-to-cell communication by transferring biomolecules within individuals or between different organisms. Trichinella spiralis is thought to secrete EVs to establish parasitism, but the detailed mechanisms of EV-mediated parasitic strategies have remained unclear. We identified EVs of T. spiralis as stage-specific in adult worms in host intestine (AW-EVs) and larvae in host muscles (ML-EVs). Stage-specific EVs exhibited unique functions, such as suppressing the expression of mucin-related genes only in AW-EVs and inducing genes associated with myoblast differentiation found in nurse cell formation in ML-EVs. Both EVs also suppressed IL-6 expression. These functions corelated with the contents of each EV, as confirmed by proteomics and whole microRNA sequencing. One of the microRNAs common to both EVs, tsp-miR-1, was involved in the suppression of IL-6 expression. These results suggest that Trichinella spiralis secretes stage-specific EVs containing unique as well as common biomolecules to regulate the host microenvironment for parasitism.

## Introduction

Extracellular vesicles (EVs) are heterogenous nano-sized vesicles secreted by cells. They participate in intercellular communication by carrying and transferring many biomolecules between cells. EVs have been found to consist of several subtypes. Generally, two main subtypes such as exosomes and microvesicles are well-defined based on their size and biogenesis. Exosomes are small EVs with a size of less than 200 nm in diameter. They are formed in the cell as intraluminal vesicles contained in multivesicular bodies by an endosomal pathway. Once the multivesicular bodies fuse with the plasma membrane, the intraluminal vesicles are released out of the cells. They are called exosomes in the extracellular compartment. Microvesicles are EVs with a size of 200-1000 nm in diameter. They are shed from the cells by budding off the plasma membrane. EVs contain many biomolecules constituted of membranes and cytosols such as lipids, proteins, and nucleic acids ^1–3^. Cumulative evidence revealed that EVs contribute to many aspects of physiological and pathogenic processes. For instance, EVs derived from ovarian cancer cells exert their function by transferring miR-205 to endothelial cells via a lipid raft-associated pathway leading to the promotion of angiogenesis *in vitro* and accelerating angiogenesis and tumor growth in the murine model ^4^.

Parasitic helminths, in general, take many complicate strategies to manipulate host responses for their long-term infections. The most obvious strategy is to deactivate the host immune defense against their infection by inactivation of the antigen-recognition system, tolerizing the immune system to their antigens, and stimulating the immuno- regulatory mechanisms. Excretory-secretory products from helminths, a mixture of many kinds of biomolecules including proteins, glycan, lipids, and nucleic acids, are most likely to take part in these functions ^5,6^. Those biomolecules are thought and investigated to be secreted from parasites in a free form. However, several studies recently revealed the EVs as a new vehicle used by helminths to manipulate host immune responses. *Heligmosomoides polygyrus*, one of the murine intestinal nematodes, secretes EVs containing small RNAs and proteins and transfers these molecules to mammalian cells. They can regulate the innate immune responses by reducing bronchoalveolar eosinophilia and suppressing the expression of type 2 cytokines, such as interleukin 5 (IL-5) and IL-13. They also suppress the expression of the IL-33 receptor on the murine intestinal cell line and the activation of bone marrow-derived macrophages ^7,8^.

*Trichinella spiralis* is a parasitic nematode that naturally infects a broad range of mammalian hosts including humans. Within the host, its life cycle consists of the intestinal and muscle stages. Following ingestion of *T. spiralis*-infected muscles, the larvae develop into adult worms in the small intestine and adult worms produce offspring which are newborn larvae. Newborn larvae then migrate into the blood circulation and deposit in the skeletal muscles, finally transform into muscle larvae waiting for ingestion of them by a new host. The intestinal stage takes about 2-3 weeks after infection, then all adult worms are expelled. In the muscle stage, newborn larvae infect myofiber, then develop into muscle larvae in parallel with transforming infected myofiber into capsule-surrounded cells (nurse cells) in which the larvae can occupy for many months to year ^9,10^. Recently, one study showed that muscle larvae of *T. spiralis* secrete EVs and speculated its biological functions in the manipulation of host microenvironments. However, whole picture and functions of EVs in *T. spiralis* infection are still obscure.

In this study, we identified EVs from two different stages of *T. spiralis* infection. We observed the specific, unique as well as common functions of each EV. Moreover, the profiling of proteins and miRNAs in each EV was also investigated. Our study may provide new insight into the understanding of parasitic mechanisms in *Trichinella* infection.

## Results

### *Trichinella spiralis*, both adult worms and muscle larvae, secrete extracellular vesicles

To characterize EVs from two different stages of *T. spiralis* such as adult worms and muscle larvae, we collected secretory products from each stage after 2-day culture in serum-free culture media. Then, adult worms-derived EVs (AW-EVs) and muscle larvae-derived EVs (ML-EVs) were purified from their secretory products by ultracentrifugation method. Freshly prepared adult worms-derived EVs (AW-EVs) and muscle larvae-derived EVs (ML-EVs) were visualized by electron microscopy. We observed EVs-like particles diameter ranging from 30 to 50 nm in both AW-EVs and ML-EVs samples as shown in Supplement Figs. 1a and 1b, respectively. The size and appearance of both EVs were corresponding to EVs secreted by *T. spiralis* from a previous report ^11^. We further analyzed EV-cell membrane fusion to confirm if the EV-like particles obtained by this isolation method were intact membrane surrounded vesicles and they were able to interact with recipient cells to operate their functions. EVs are surrounded by lipid-enriched membrane structures like cell membranes. With the high lipid composition of EVs and cell membranes, they are tended to interact with each other by membrane fusion ^2,3,12^. In order to examine EV-cell membrane fusion, ML-EVs were labeled by staining with carboxyfluorescein succinimidyl ester (CFSE), a membrane-permeable fluorescent dye, before co-incubation with HT-29 cells (human colon cancer cell line). Thirty minutes after co-incubation of CFSE-labeled EVs and HT-29 cells, we performed flow cytometry analysis to detect a signal of CSFE in HT-29 cells. We found that HT-29 cells that were co-incubated with CFSE-labeled EVs showed higher mean fluorescence intensity (MFI) of CFSE than cells that were co-incubated with CSFE dye alone (negative control of CFSE-labeled EVs: Phosphate buffered saline (PBS) that were stained by CFSE with concentration and washing steps same as staining of EVs by CFSE) (Supplementary Fig. 1c), indicating the fusion of CFSE-labeled EVs and cell membrane of HT-29 cells. These results suggested that the EV-like particles obtained by this isolation method were intact vesicles capable of interacting with recipient cells. Thus, this isolation method was used for isolating EVs for further functional analysis of EVs.

### AW-EVs and ML-EVs play a common role on suppression of macrophage activation

In order to investigate the common function of both EVs, we therefore tested the influence of EVs on the classical macrophage activation induced by lipopolysaccharide (LPS). RAW 264.7 cells, a murine macrophage cell line, were prior co-cultured with AW-EVs or ML-EVs for 24 hours then the cells were stimulated by LPS. Six hours after LPS stimulation, macrophage activation markers such as a pro-inflammatory cytokine, IL-6, and a costimulatory molecule, CD80, were examined. Quantitative reverse transcription polymerase chain reaction (RT-qPCR) revealed that *Il6* mRNA was up-regulated in the cells that stimulated by LPS but not in unstimulated cells. Moreover, among LPS-stimulated cells, the cells that were prior co-cultured with AW-EVs or ML-EVs exhibited a lower level of *Il6* mRNA compared to LPS-stimulated control cells (Fig. 1a), indicating that AW-EVs and ML-EVs suppressed LPS-induced upregulation of *Il6* mRNA in RAW 264.7 cells. The suppressive effect of both EVs also extended to LPS-induced secretion of IL-6 from RAW 264.7 cell in dose-dependent manners (Fig. 1b). In addition, both EVs also markedly inhibited LPS-induced CD80 expression on the cell surface of RAW 264.7 cells in dose-dependent manners (Fig. 1c). These results suggested that the two difference stages of *T. spiralis* which survive in very different physiological conditions secrete EVs to play a common role in the suppression of macrophage activation.

**Fig. 1.**
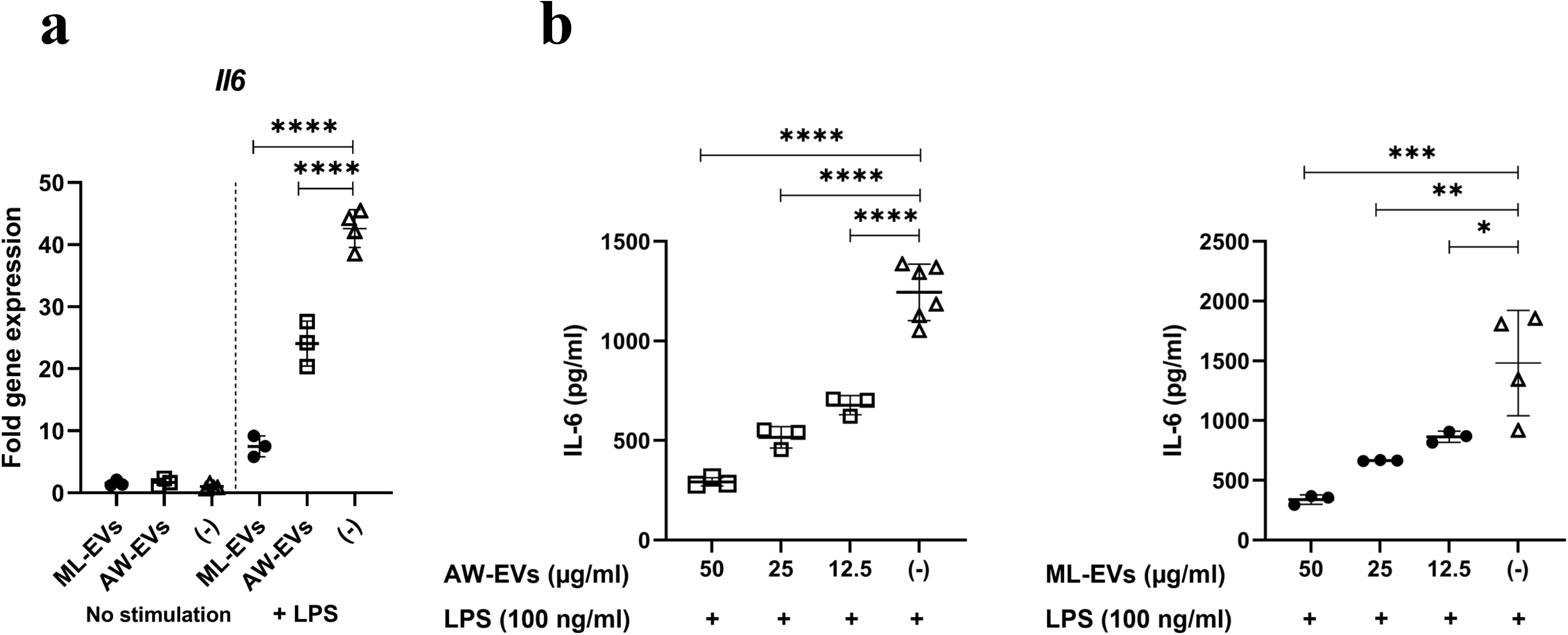
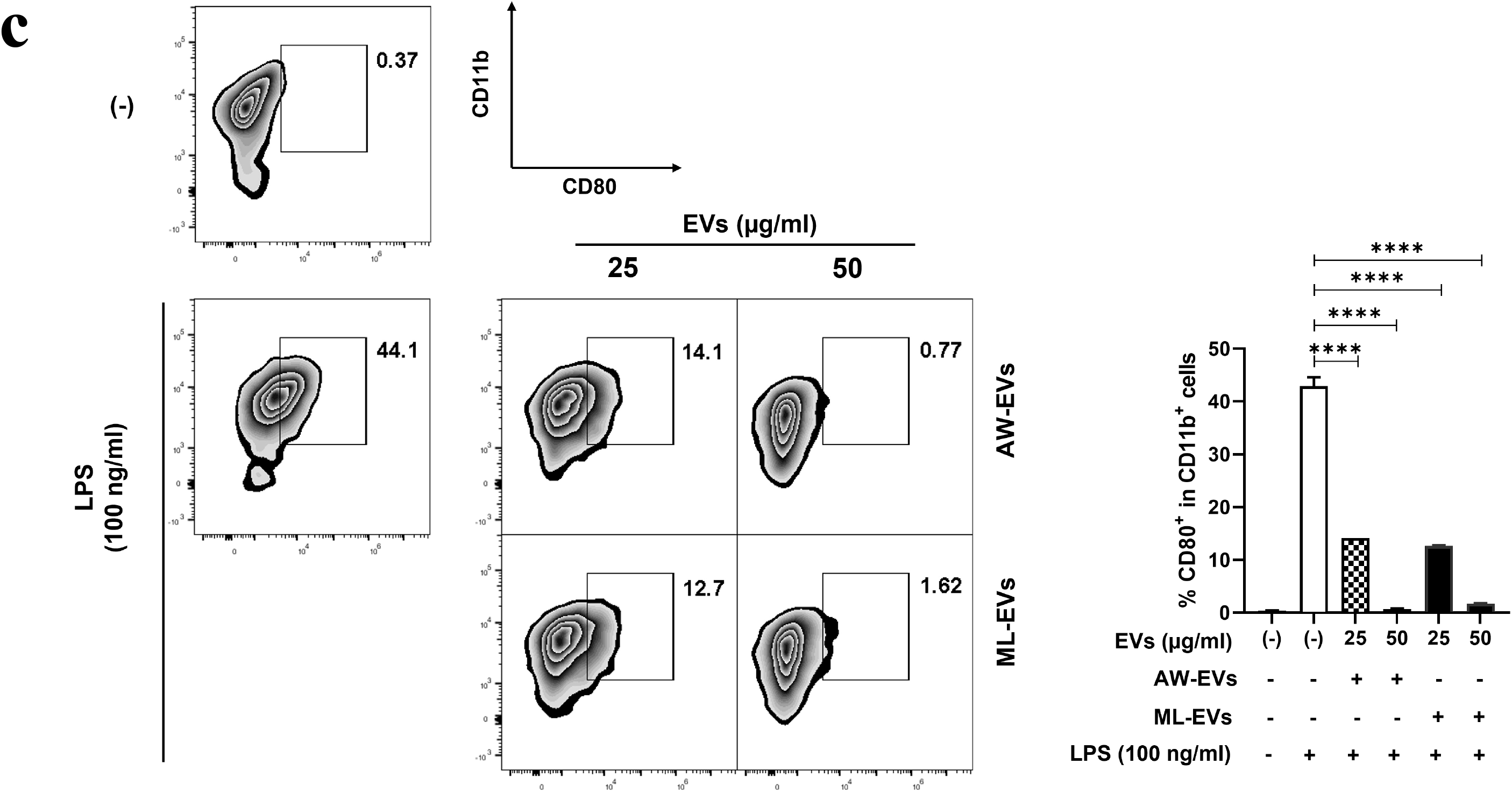
AW-EVs and ML-EVs suppress expression of macrophage activation markers. Expression of macrophage activation markers in RAW 264.7 cells including *Il6* mRNA, IL-6 released in culture supernatant and CD80 were examined after 24 hours pre-treatment with AW-EVs, ML-EVs or PBS followed by 6 hours stimulation with 100 ng/ml of LPS. **(a)** Relative expression of *Il6* mRNA in RAW 264.7 cells that were co-cultured with 12.5 µg/ml AW-EVs (AW-EVs), 12.5 µg/ml ML-EVs (ML-EVs) or PBS ((−)) prior to stimulated by LPS (+LPS) or without stimulation (No stimulation). The normalized *Il6* mRNA of each sample is relative to normalized *Il6* mRNA of PBS-treated cells ((−)) without LPS stimulation (No stimulation). The relative expression is presented as fold gene expression. Data are representative of three independent experiments. **(b)** The levels of IL-6 in culture supernatant of RAW 264.7 cells that were co-cultured with various concentration of AW-EVs or ML-EVs or PBS prior to stimulated by LPS. The levels of IL-6 were measured by Enzyme-linked immunosorbent assay (ELISA). Data are representative of at least three independent experiments. **(c)** Representatives and frequencies of CD80-expressing RAW 264.7 cells (CD80^+^CD11b^+^cells) that were co-cultured with 25 or 50 µg/ml of AW-EVs or ML-EVs or PBS ((−)) prior to stimulated by LPS. CD80 expression was measured by flow cytometry. Data are representative of three independent experiments. Each dot represents an individual biological replicate. Error bars denote mean value ± SD (**A** and **B**). Graph presents mean values ± SD (n=2) (**C**). *, *p*<0.05. **, *p*<0.01. ***, *p*<0.001. ****, *p*<0.0001. The statistically significant were calculated using one-way ANOVA.

### AW-EVs play a specific role in the suppression of mucin-related gene expression

An increase of mucin production at the intestinal epithelium is an important mechanism to enhance worm expulsion ^13^. We, therefore, hypothesized that adult worms of *T. spiralis* which reside in a small intestine may secrete EVs with a specific function to suppress mucin production. Reducing mucin production would cause a delay in worm expulsion, which prolongs the survival time of *T. spiralis* adult worms in the intestine, giving benefits to them in which they have enough time and an appropriate microenvironment for their development and producing offspring. In order to examine the specific function of AW-EVs, HT-29 cells, a human colorectal adenocarcinoma cell line, were co-cultured with AW-EVs, ML-EVs or PBS. Then mRNA expression of mucin-related genes was examined by RT-qPCR. We found that only AW-EVs but not ML-EVs could significantly down-regulate mRNA expression of *MUC1* and *MUC5AC* in HT-29 cells as shown in Figs. 2a and 2b, respectively. Next we confirmed the specific function of AW-EVs in more physiological condition of small intestine using small intestinal organoids. From this experiment, unfortunately we failed to detect the expression of *Muc1* and *Muc5ac* mRNAs in the organoids. We then focused on other mucin-related genes that have been reported to have critical effects on intestinal helminth infection i.e. Trefoil factor 3 (*Tff3*) and Chloride channel accessory 3 (*Clca3*) ^14,15^. Twenty-four hours after treating the organoids with AW-EVs, ML-EVs or PBS, we found that only AW-EVs significantly down-regulated expression of *Tff3* mRNAs but not ML-EVs- or PBS (Fig. 2c). AW-EVs also down-regulated the expression of *Clca3* mRNAs (Fig 2d). These results indicated that AW-EVs play a specific function in the suppression of mucin-related gene expression in intestinal epithelium cells.

**Fig. 2.**
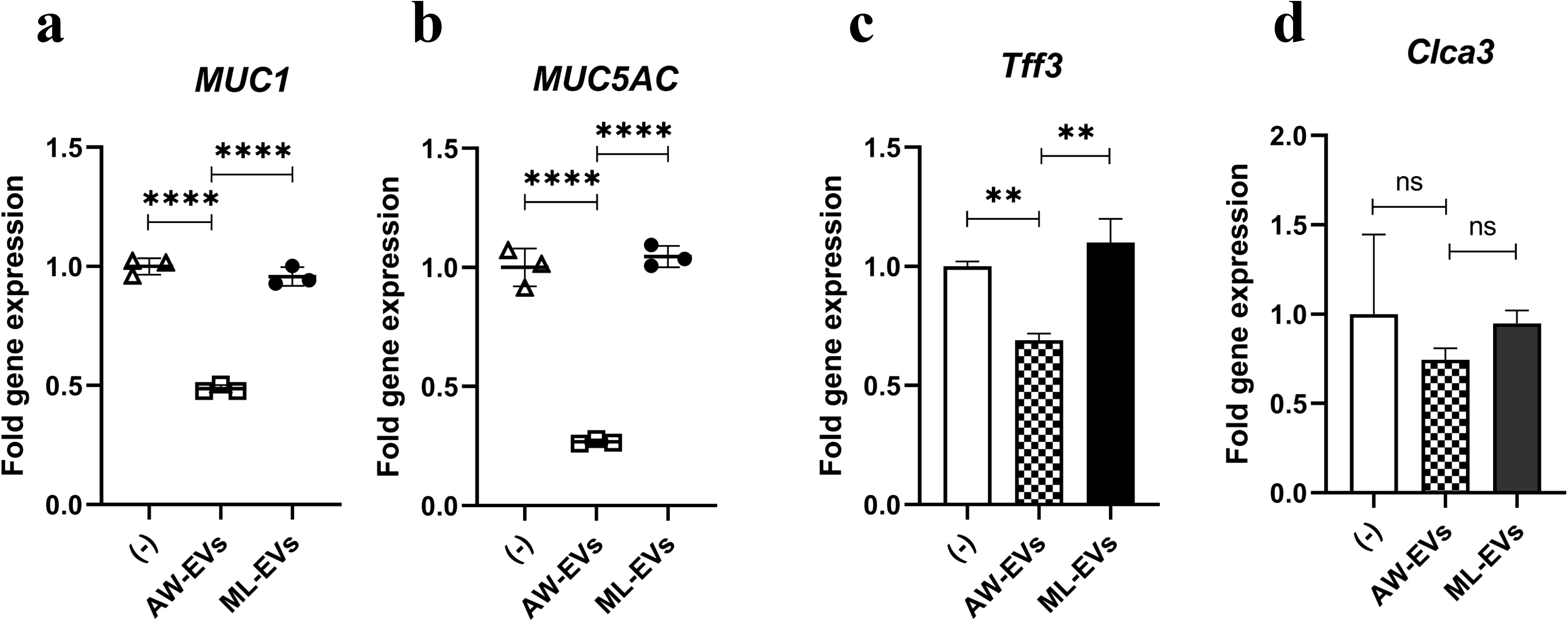
AW-EVs suppress mucin related genes in HT-29 cells and small intestinal organoids. Expression of mucin genes in HT-29 cells and mucin related genes in mouse intestinal organoids were measured by RT-qPCR 24 hours after treatment with 12.5 µg/ml AW-EVs, 12.5 µug/ml ML-EVs or PBS. **(a** and **b)** Relative expression of *MUC1* **(a)** and *MUC5AC* **(b)** in HT-29 cells that were treated with AW-EVs, ML-EVs or PBS ((−)). Data are representative of three independent experiments. **(c** and **d)** Relative expression of *Tff3* and *Clca3* in mouse intestinal organoids that were treated with AW-EVs, ML-EVs or PBS ((−)). Data are representative of three independent experiments. The normalized target mRNAs of each sample are relative to normalized target mRNAs of PBS-treated cells ((−)) (**a** and **b**) or PBS-treated organoids ((−)) (**c** and **d**). The relative expressions are presented as fold gene expression. Each dot represents an individual biological replicate. Error bars denote mean value ± SD (**a** and **b**). Graphs present mean values ± SD (n=2) (**c** and **d**). ns, not significant. **, *p*<0.01. ****, *p*<0.0001. The statistically significant were calculated using one-way ANOVA.

### ML-EVs play a specific role in the enhancement of myoblast differentiation

Next, we investigated a specific function of ML-EVs by focusing on myoblast differentiation, which is one of many processes that contribute to nurse cell formation. myoblast differentiation is triggered once the myofiber gets the infection with a newborn larva of *Trichinella*. Then myoblast (satellite cell) which resides on myofiber is activated to proliferate and differentiate to incorporate into Trichinella-infected myofibers to form the nurse cells ^16^. In order to test the specific function of ML-EVs on myoblast differentiation, we cultured C2C12 cells, a murine myoblast cell line in differentiation media to induce the differentiation as well as to co-cultured them with AW-EVs, ML-EVs, or PBS for 7 days. Then we examined the mRNA expression of genes related to myoblast differentiation including myogenin (*MyoG*) and myosin heavy chain embryonic (*Myh3*) ^17–20^. Four days after induction of the differentiation, we observed that the expression of *Myog* mRNA in the cells that were cultured in the differentiation media was up-regulated but not in the cells that were cultured in the normal growth media. Among the cells that were cultured in the differentiation media, ML-EVs-treated cells showed significantly higher expression of *Myog* mRNA than AW-EVs-treated and PBS-treated cells (Fig. 3a). In addition, 7 days after the differentiation, expression of *Myh3* mRNA in ML-EVs-treated cells also showed a significantly greater level than in other cells (Fig. 3b). These data indicated that only ML-EVs but not AW-EVs could affect the C2C12 cell differentiation induced by the differentiation media. We further confirmed the function of ML-EVs *in vivo* by direct injection of EVs into striated muscle of mice. Corresponding to the *in vitro* C2C12 cell differentiation, we found that ML-EVs caused significantly higher expression of *Myog* and Myogenic Regulatory Factor 4 (*Mrf4*) mRNAs than AW-EVs and PBS in mice striated muscle at 2 days after the injection (Figs. 3c and 3d). The contribution of *Myog* in *T. spiralis* infection during nurse cell formation has been reported ^21^. However, the involvement of *Mrf4* was unclear and *Myh3* was unknown, so we confirmed the contribution of *Mrf4* and *Myh3* in *T. spiralis-*infected mice. We found that all three genes were gradually upregulated from the first until the third week of the infection which was the initiation period of nurse cell formation (Supplementary Fig. 2). These results suggested that ML-EVs might play specific roles in the nurse cell formation via enhancement of myoblast differentiation.

**Fig. 3.**
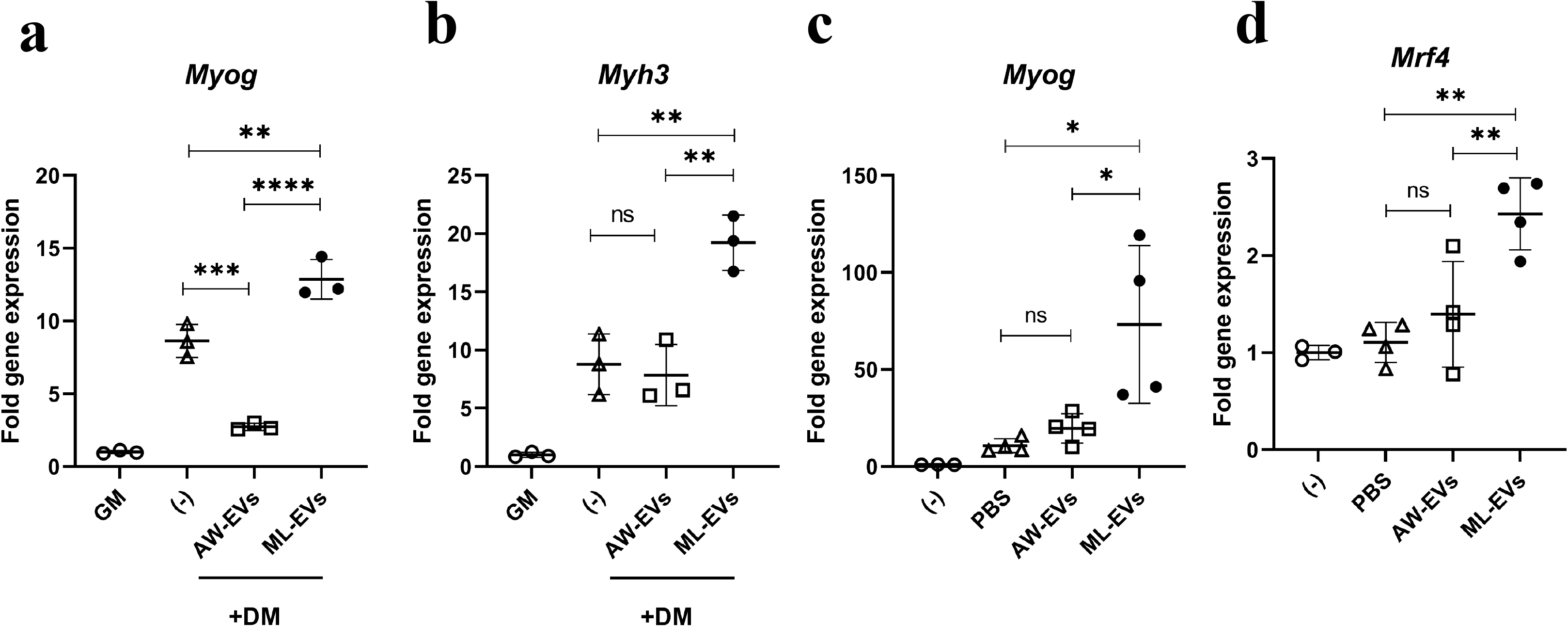
ML-EVs enhance differentiation of myoblast in vitro and in vivo. Expression of genes related to differentiation of myoblast in C2C12 cells and mouse tibialis anterior muscle were measured by RT-qPCR after treatment with AW-EVs, ML-EVs or PBS **(a** and **b)** Relative expression of *Myog* **(a)** and *Myh3* **(b)** in C2C12 cell lines 4 and 7 days respectively after cultured under normal growth media (GM) or co-cultured with 2.5 µg/ml AW-EVs, 12.5 µg/ml ML-EVs or PBS under differentiation media (DM). The normalized target mRNAs in each sample are relative to normalized target mRNAs of cells that culture in GM. The relative expressions are presented as fold gene expression. Data are representative of three independent experiments. **(a** and **b)** Relative expression of *Myog* **(c)** and *Mrf4* **(d)** in mouse tibialis anterior muscle 2 days after injection by 7.5 µg AW-EVs, 7.5 µg ML-EVs or PBS. The normalized target mRNAs of each sample are relative to normalized target mRNAs of un-injected tibia anterior muscle ((−)). The relative expressions are presented as fold gene expression. Each dot represents an individual biological replicate (**a** and **b**) or an individual mouse (**c** and **d**). Error bars denote mean value ± SD. *, *p*<0.05. **, *p*<0.01. ***, *p*<0.001. ****, *p*<0.0001. The statistically significant were calculated using one-way ANOVA.

### Proteomics reveals AW-EVs and ML-EVs contain common and specific proteins and predicted the function of EV protein

EV functions depend on the biomolecular contents in them. We then investigated the protein contents of both EVs to see whether the common function and specific function of each EVs are based on their contents. In order to identify protein contents in both EVs, we conducted liquid chromatography-electrospray tandem mass spectrometry (LC-MS/MS) analysis of AW-EVs and ML-EVs. From both EVs, we identified total of 322 proteins, 260 proteins from AW-EVs and 230 from ML-EVs. Protein profiling of both EVs showed that each EVs contained common and stage-specific ones as shown in Fig. 4a. One hundred and sixty-eight equivalent proteins were identified in both EVs, which they were thought common EV proteins. On the other hand, 92 and 62 proteins were specifically identified in AW-EVs and ML-EVs, respectively. They were thought to be AW-EV specific and ML-EV specific proteins, respectively. The list of representatives of common EV proteins as well as top ten most abundant AW-EV specific and ML-EV specific proteins are shown in Supplementary Table 1. These data suggested that the common as well as the stage specific functions of both EVs may base at least in part on their protein contents.

**Fig. 4.**
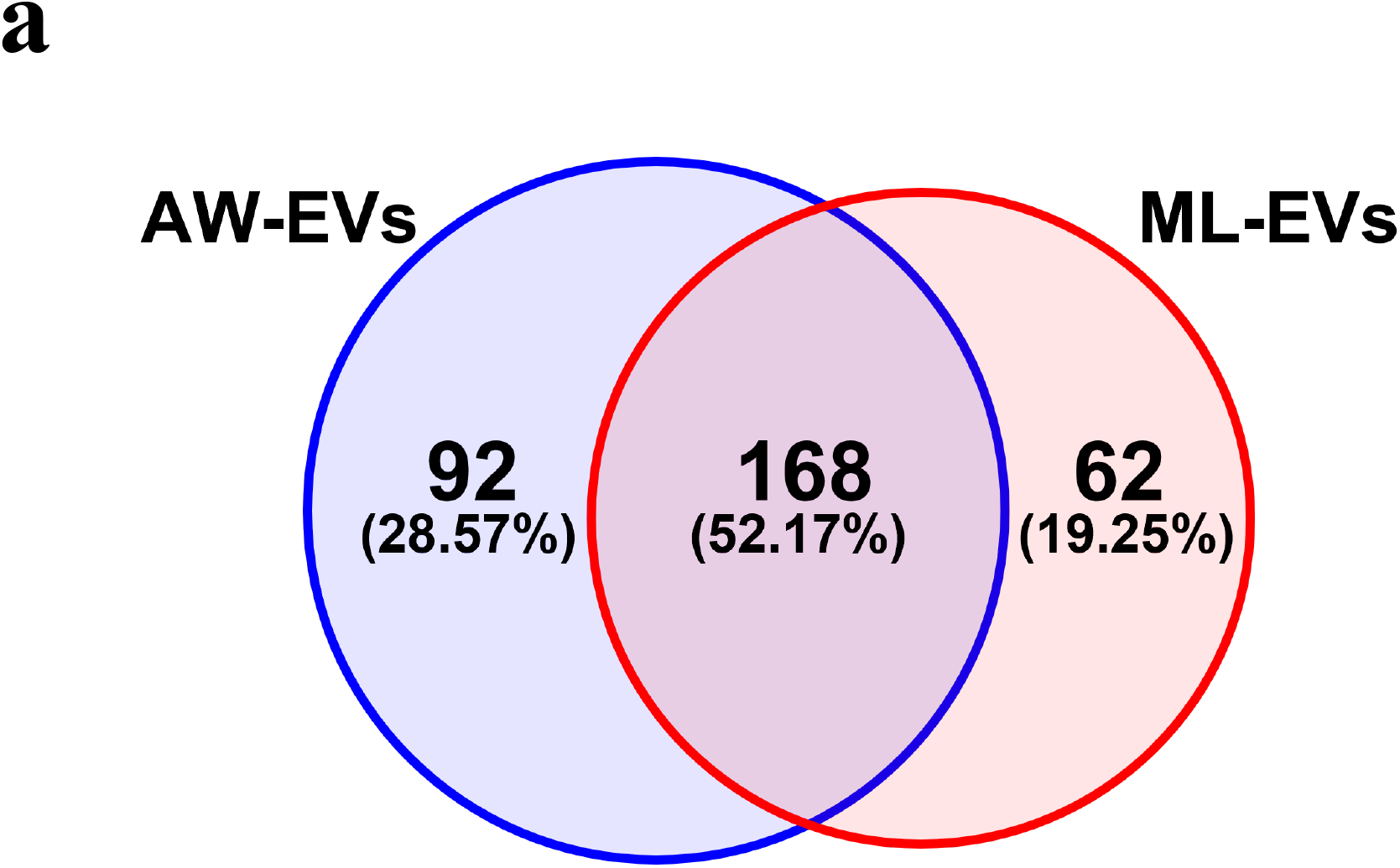
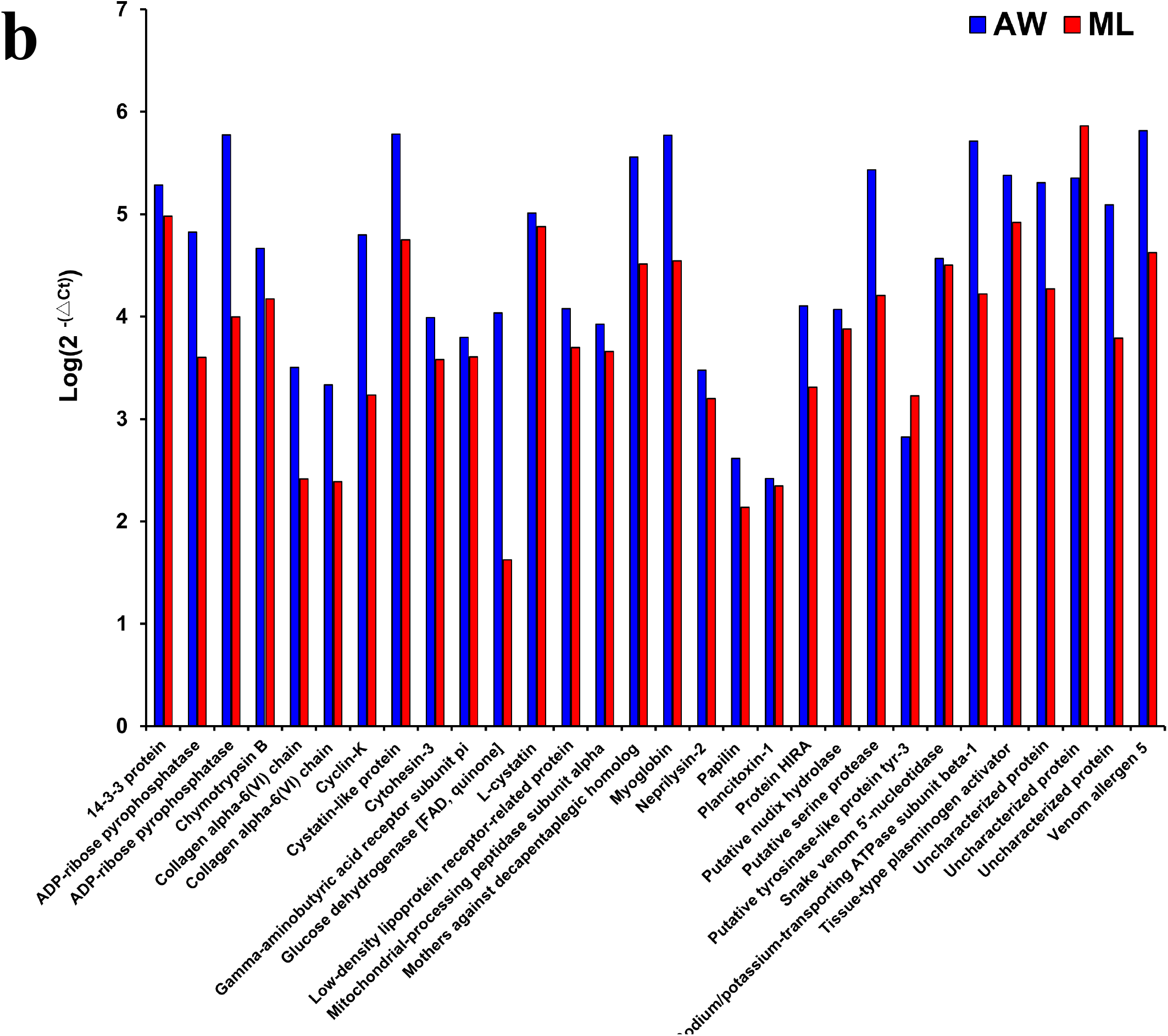
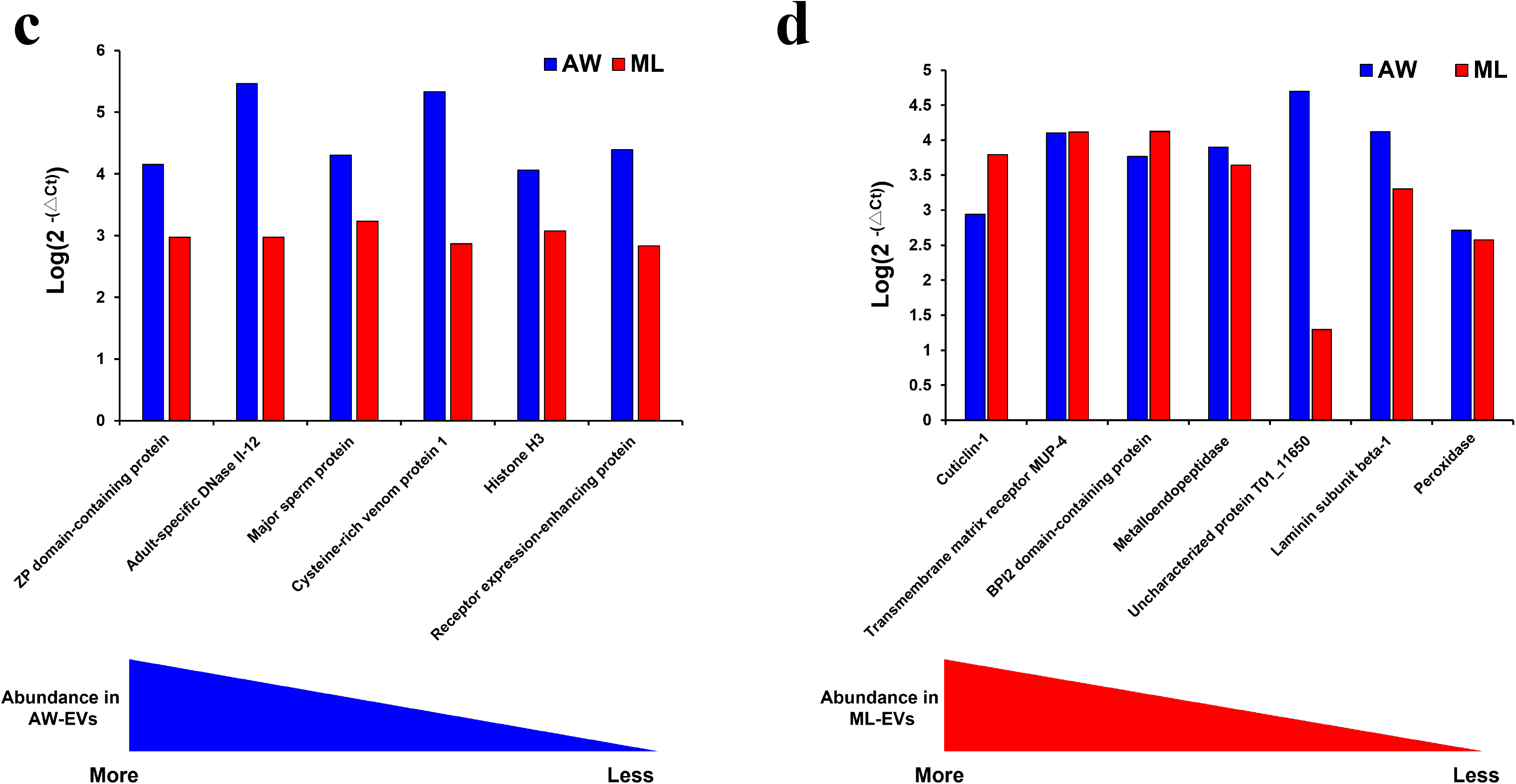
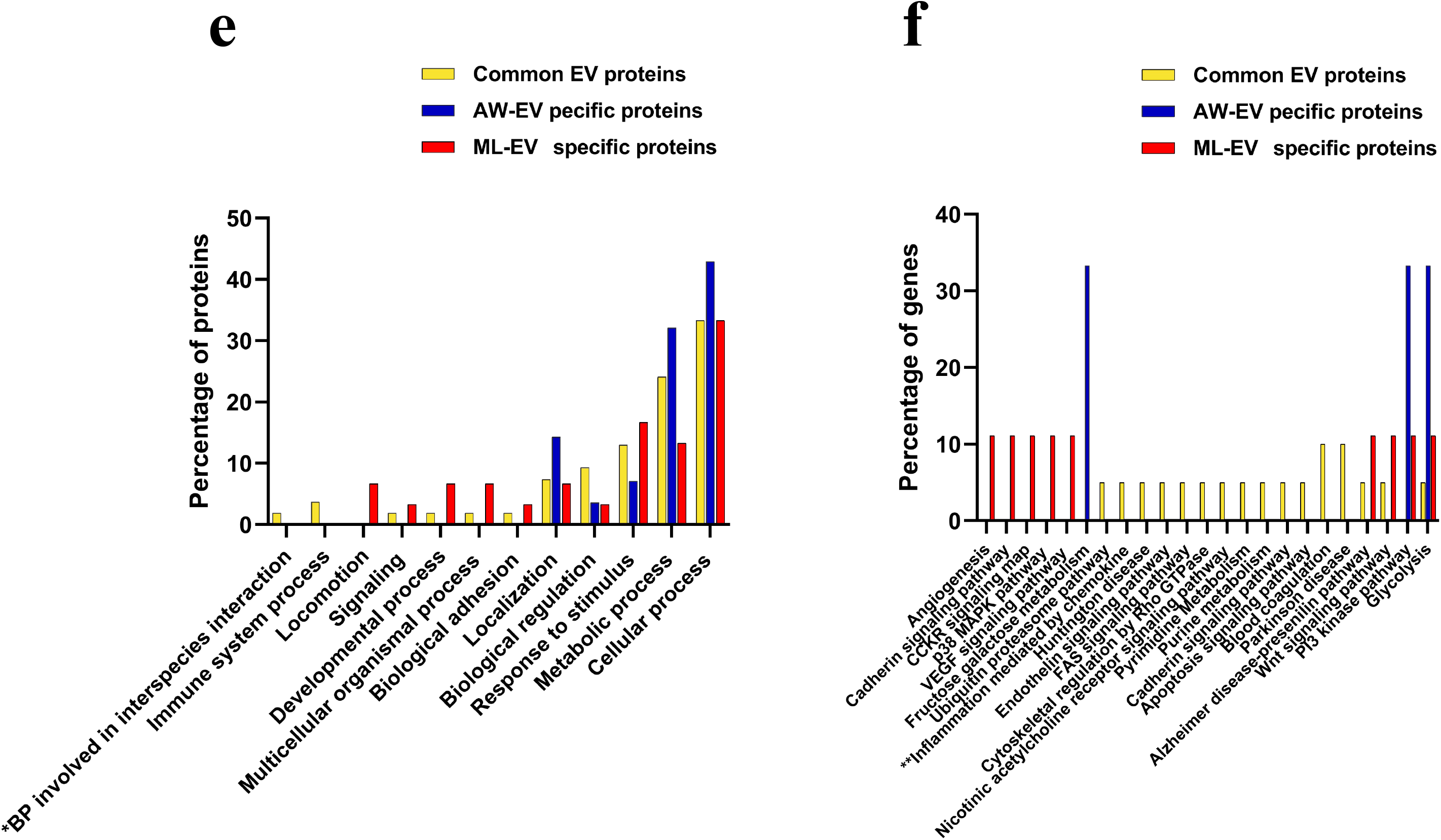
Proteomics of AW-EVs and ML-EVs. (**a**) Distribution of proteins identified in AW-EVs and ML-EVs by LC-MS/MS analysis. Venn diagram shows the number and percentage of common EV, AW-EV specific and ML-EV specific proteins. The proteomics experiment was performed in three biological replicates. (**b-d**) Expression of mRNA of EV proteins in worm bodies. Expression of mRNA of common EV (**b**), AD-EV specific (**c**) and ML-EV specific (**d**) proteins in adult worms (AW) and muscle larvae (ML) were analyzed by RT-qPCR. Blue or red triangle illustrate the order of individual proteins in bar graph based on their abundance (Quantitative value) in EVs. The abundance was order from high to low (**c** and **d**). The mRNAs expression level are presented as Log(2 ^-(△Ct)^). The detail of calculations is explained in Materials and Methods. (**e** and **f**) Functional analysis of EV proteins based on Gene ontology. Gene ontology analysis of common EV, AD-EV specific and ML-EV specific proteins were analyzed by The PANTHER classification system in term of biological function **(e)** and Panther pathway **(f)**. **\***, Full name is biological process involved in interspecies interactions between organisms. **\*\***, Full name is Inflammation mediated by chemokine and cytokine signaling pathway.

We further examined the expression of mRNAs of EV proteins in adult worms and muscle larvae to investigate whether the stage-specific EV proteins are expressed in the stage-specific manner. RT-qPCR analysis revealed that both adult worms and muscle larvae expressed mRNAs of common EV proteins as we expected (Fig. 4b). However, the mRNAs of stage-specific EV proteins were not specifically expressed in the stage-specific manner, but they were expressed in both stages of *T. spiralis*. AW-EV specific proteins were supposed to be expressed only in adult worms, but muscle larvae also expressed AW-EV specific proteins (Fig. 4c). Similarly, ML-EV specific proteins were expected to be expressed in muscle larvae only but adult worms also expressed ML-EV specific proteins (Fig 4d), indicating that *T. spiralis* might regulate the release of stage-specific EVs and/or packing of specific EV proteins into EVs in the stage-specific manner. We had the limitation to identifying the function of individual EV proteins by wet lab experiments. So, we performed a computer-based functional analysis of EV proteins using Genes Ontology Analysis to determine what kind of biological processes or pathways are implicated by EV proteins and whether those biological processes or pathways contribute to the establishment of parasitism by *T. spiralis*. We separately analyzed sets of EV proteins according to their stage-specific groups such as common EV, AW-EV specific and ML-EV specific proteins. We found that common EV proteins are involved in many biological processes that were also shared with AW-EV or ML-EV specific proteins. However, the most interesting processes exclusively seen in the common EV proteins were ‘biological process involved in interspecies interactions between organisms’ and ‘immune system process’ (Fig. 4e). From this result, we speculated that both adult worms and muscle larvae of *T. spiralis* secrete the same set of proteins via EVs to interact with hosts and to regulate the host immune system for initiating parasitism. Some ML-EV specific proteins were preferentially involved in several pathways especially angiogenesis and VEGF signaling pathway (Fig. 4f). These two pathways are processes that might contribute to forming new blood vessels with highly permeable property on outer surface of collagen capsule of nurse cells to provide nutrients for muscle larvae and to discard waste products from them ^22,23^ again suggesting that muscle larvae secrete EVs containing specific proteins promoting nurse cell formation. For AW-EV specific proteins, on the other hand, they uniquely implicate in fructose galactose metabolism, but how much these pathways are important to *T. spiralis* infection is unknown.

In addition, EV protein markers and EV-associated proteins such as heat shock protein 70 (HSP70), 14-3-3 protein, and actin-5C, which have been found in EVs from other organisms including helminths, were also identified in both EVs samples (Supplementary Table 2). This could confirm that the EV-like particle that we identified by electron microscopy were real EVs. Moreover, some proteins in EVs from *T. spiralis* were conserved in other helminths.

### AW-EVs and ML-EVs contain common and specific miRNAs and EV miRNAs target genes related to EV function

Extracellular vesicles contain microRNA (miRNA) also in addition to the proteins. We performed whole miRNAs sequencing analysis to identify miRNAs in both EVs. The result was similar to that in protein contents, in which both EVs contained common and stage-specific miRNAs. From total of 175 miRNAs identified in both AW-EVs and ML-EVs, 103 miRNAs matched with known miRNAs in miRBase (known miRNAs) and 72 did not match with any miRNAs in miRBase (unknown miRNAs). AW-EVs contained 132 miRNAs. ML-EVs contained 97 miRNAs. Fifty-four miRNAs were common miRNAs between the two EVs. Seventy-eight and 43 were AW-EV specific- and ML-EV specific-miRNAs, respectively, as shown in Fig. 5a. Representatives of known common- and top ten most abundant AW-EV specific- and ML-EV specific-miRNAs were shown in Supplementary Table 3. We then examined the expression of EV miRNAs in worm bodies. The result was similar to the expression profiles of mRNA of EV proteins. Common EV-, AW-EV specific- and ML-EV specific-miRNAs were expressed in both adult worms and muscle larvae (Figs. 5b-5d).

**Fig. 5.**
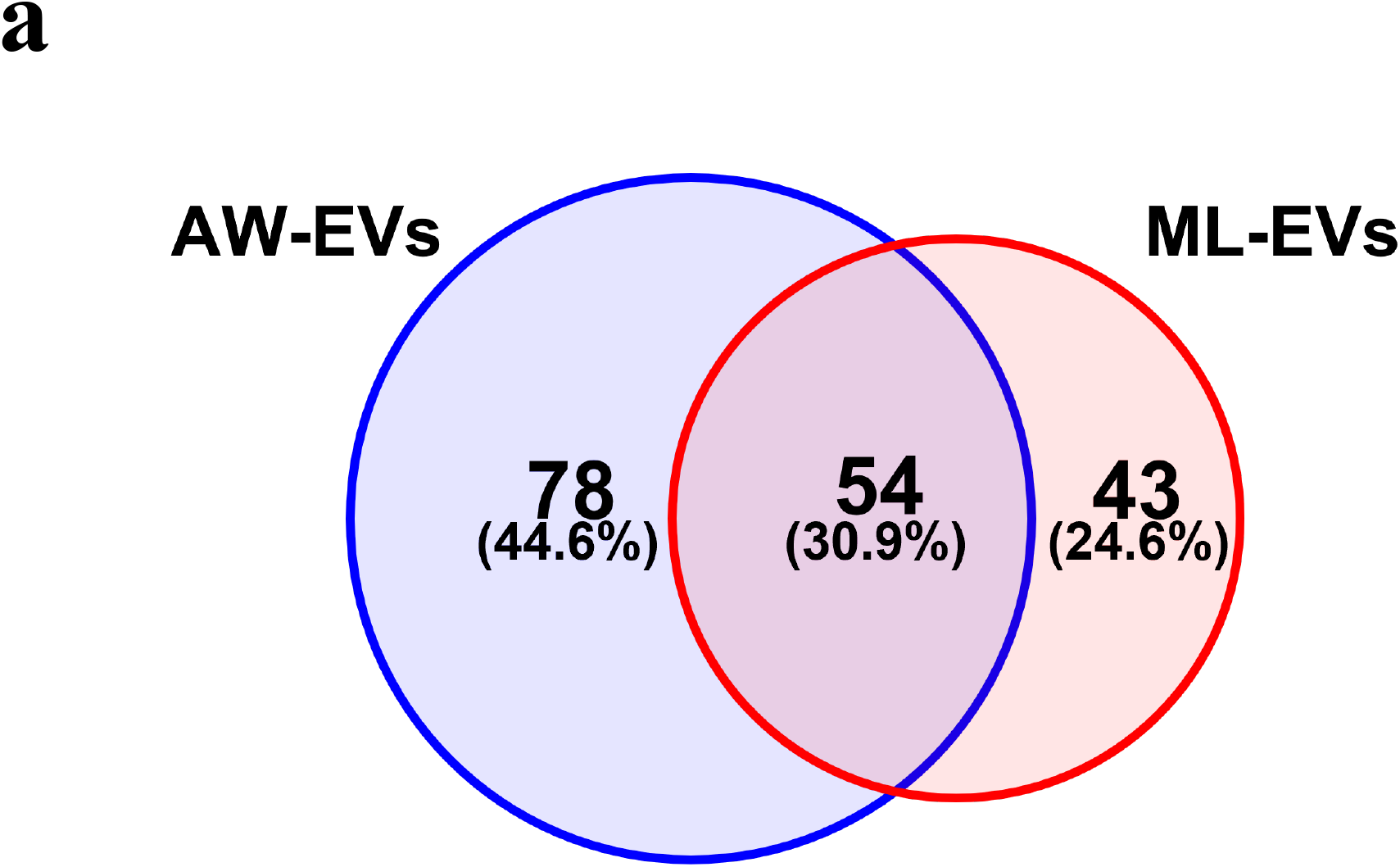
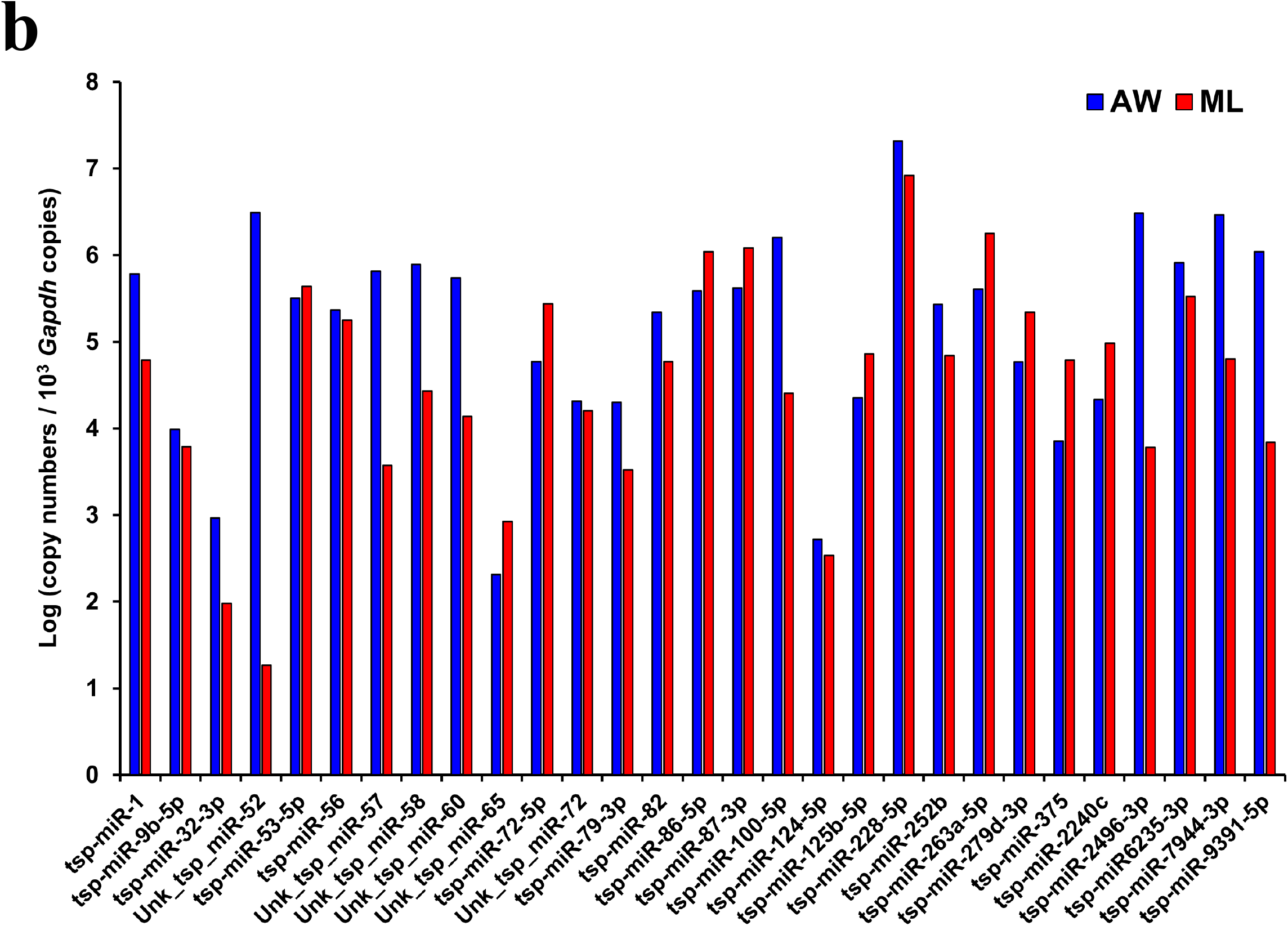
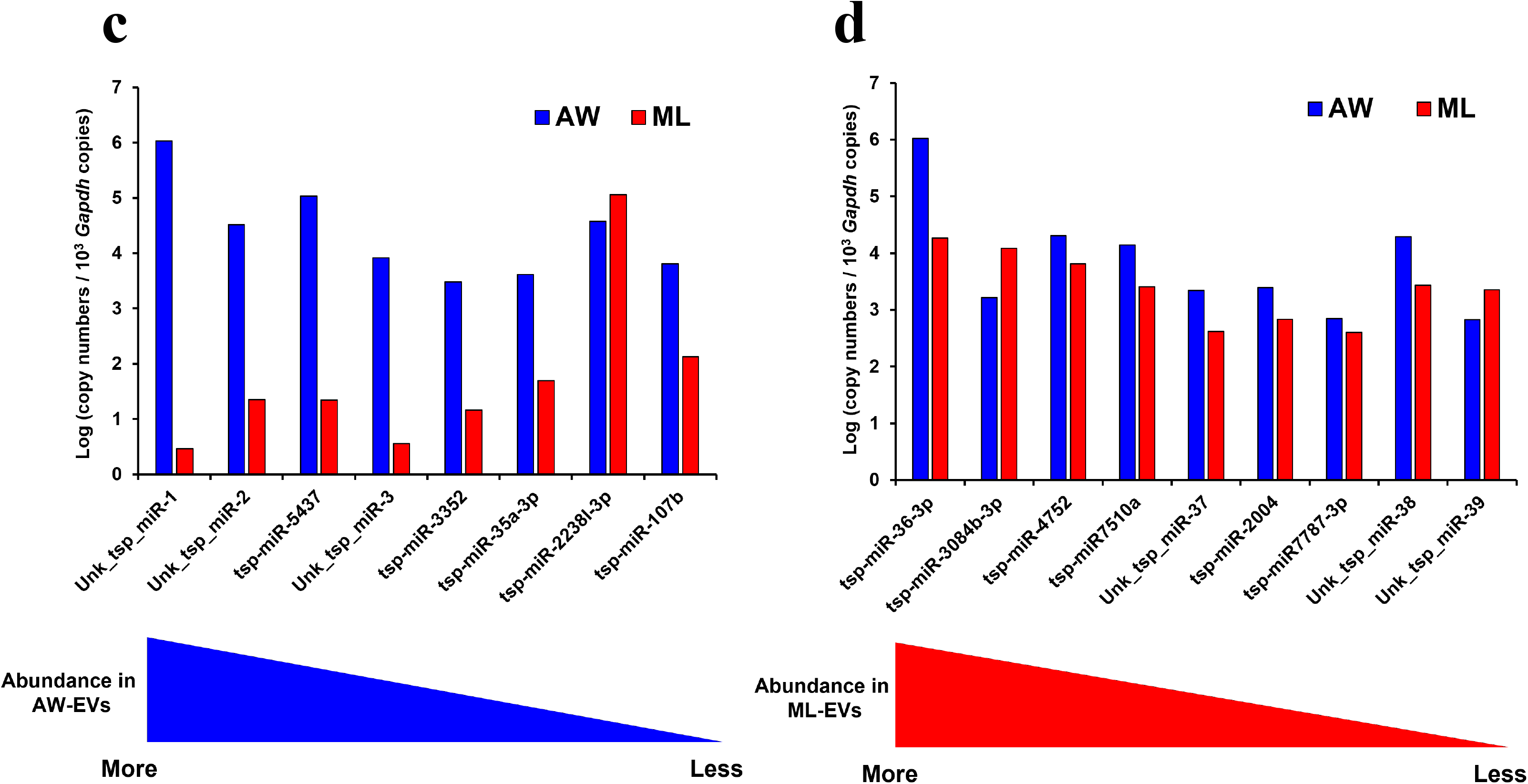
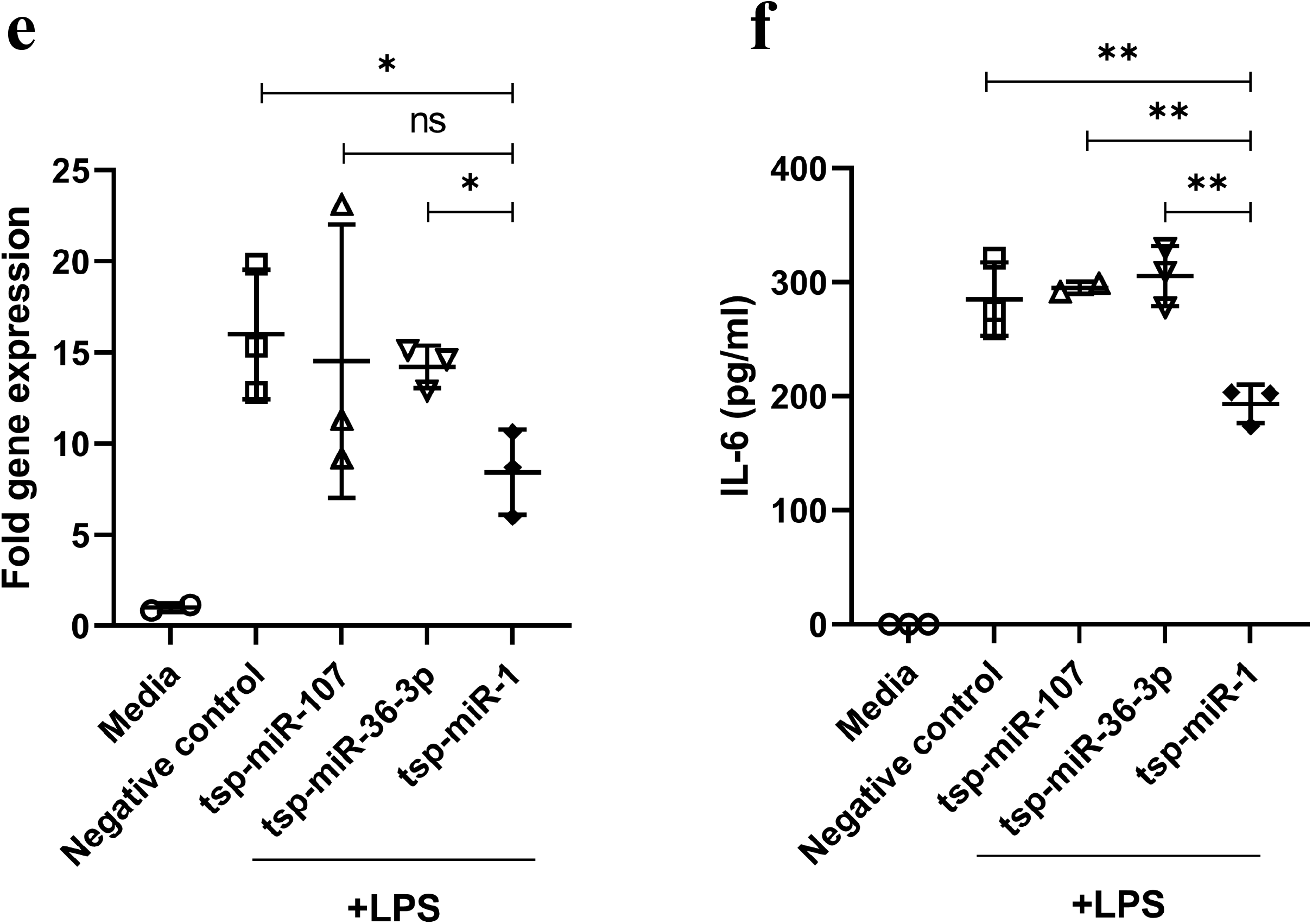
Profiling of miRNAs identified in AD-EVs and ML-EVs and effect of synthetic EV miRNAs on macrophage activation. (**a**) Distribution of miRNAs identified in AW-EVs and ML-EVs by miRNA-sequencing analysis. Venn diagram shows number and percentage of common EV, AW-EV specific and ML-EV specific miRNAs. (**b-d**) Expression of common EV (**b**), AD-EV specific (**c**) and ML-EV specific (**d**) miRNAs in adult worms (AW) and muscle larvae (ML) were analyzed by RT-qPCR. Blue or red triangle illustrate the order of individual miRNA in bar graph based on their abundance (Normalized read counts) in EVs. The abundance was order from high to low (**c** and **d**). (**b**) Relative expression of *Il6* in RAW 264.7 cells that were transfected with assigned synthetic miRNAs prior to stimulated by LPS. Data are representative of two independent experiments. The normalized *Il6* mRNA of each sample is relative to normalized *Il6* mRNA of un-transfected cells that culture without LPS stimulation ((−)). The relative expression is presented as fold gene expression. (**c**) The levels of IL-6 in culture supernatant of RAW 264.7 cells that were transfected with assigned synthetic miRNAs prior to stimulated by LPS. The levels of IL-6 were measured by ELISA. Data are representative of two independent experiments. Each dot represents an individual biological replicate. Error bars denote mean value ± SD (**b** and **c**). ns, not significant. *, *p*<0.05. **, *p*<0.01. The statistically significant were calculated using one-way ANOVA (**b-c**).

Next, we investigated the possibility that miRNAs contained in each EVs act as biomolecules that regulate the common- and stage-specific function of AW-EVs and ML-EVs. We performed the prediction of miRNA-target genes, which meet the functional analysis experiments, including *Il6*, *MUC5AC* and *Myog*. We found that murine *Il6* was one of the target genes of tsp-miR-1, which was contained in both EVs. *MUC5AC* was predicted to be one of the target genes of AW-EV specific tsp-miR-107. *Mstn*, which have been reported to regulate the expression of *Myog* during differentiation of myoblasts ^24^ was predicted as one of the target genes of ML-EV specific tsp-miR-181b-5p. In order to validate the function of each miRNAs on their predicted target genes, tsp-miR-1, tsp-miR-107 and tsp-miR-181b-5p were synthesized and tested. Two more miRNAs were selected to be synthesized because tsp-miR-5437 was the most abandant in AW-EVs and tsp-miR-36-3p was also in ML-EVs specific miRNAs. We transtrasfected the synthesyzed miRNAs to RAWs 264.7, HT-29 and C2C12 cells. Twenty-four hours after transfection of synthesized miRNAs into RAW 264.7 cells followed by 6 hours stimulation with LPS, we found that only tsp-miR-1 down-regulated the expression of *Il6* mRNA and protein as shown in Figs. 5e and 5f, respectively. On the other hand, we did not see any effect of tsp-miR-107 or tsp-miR-5437 on the expression of *MUC5AC* in HT-29 cell lines (data not shown). For miR-36-3p and miR-181b-5p, our experiment did not show that they had any effect on *Myog* expression during the C2C12 cell differentiation (data not shown). These results suggested that tsp-miR-1 might act as a main biomolecule to suppress the expression of IL-6 in the mouse.

## Discussion

To establish long-term infection, helminth parasites use many strategies to altering host cell response. The mechanism underlying these strategies have not been fully understood. In this study, we aimed to clarify the use of EVs by *T. spiralis* as a mechanism for manipulating host microenvironment. We illustrated the common and different functions as well as biomolecular contents in EVs from two different stages of *T. spiralis*. We found that *T. spiralis* used EVs as carriers of biomolecules which contribute to altering host responses for optimizing favorable conditions for the parasite to take a long period of infection with less harm to the host.

We showed that EVs from both infectious stages of *T. spiralis* suppressed the activation of inflammatory macrophages. Previous reports illustrated that the infection by *T. spiralis* ameliorates several inflammatory diseases such as type 1 diabetes, inflammatory bowel disease, allergic airway inflammation, and autoimmune encephalitis. Some of the ameliorative effects caused by *T. spiralis* infection are due to the bias to elicit type 2 immune responses ^25–28^. There are also many studies regarding the parasite-derived molecule(s) which suppress the inflammatory macrophage activation and instead induce type 2 response including alternatively activated macrophages. However, the detailed molecular mechanisms underlying the biological actions of most of them are unclear. In general, the expulsion of helminth by the host is thought to depend on type 2 immune responses such as IgE production and eosinophil/basophil activations and so on. According to this aspect, the host not parasites might actively induce the type 2 response. Indeed, Chitin, a main component of helminth outer skin, is reported to induce type 2 response via sensing by host chitinase-relating pathway ^29^, indicating the active induction of type 2 response by the host. However, in this study, we demonstrated that tsp-miR-1, the one component in *T. spiralis*-derived EVs, suppressed the IL-6 production from the macrophage cell line, suggesting *T. spiralis* might actively suppress the classical inflammatory response. Since the classical inflammatory response including activation of M1 macrophages is associated with the expulsion of larvae in the muscle ^30,31^, it is reasonable that *T. spiralis* actively suppresses IL-6 production at least in the muscle larvae stage. On the other hand, it is unknown the reason why adult worms in the intestine secrete tsp-miR-1 capable of suppressing IL-6 production. It might be possible that tsp-miR-1 may suppress excessive destruction of the intestinal tissue by the classical inflammatory response to establish long-term attachment of adult worms to the intestinal walls. Alternatively, tsp-miR-1 might modify the expression of other target genes except for *Il6* for their parasitism. Further investigation is needed to clarify this issue.

The expulsion of helminth in the intestinal tract is largely dependent on mucin production from the host goblet cells. Prolonged expulsion of adult worms from the intestine was seen in *Muc5a*-deficient mice infected with *T. spiralis* ^32^, indicating the importance of mucin in the expulsion of adult worms of *T. spiralis* from the intestine. In this study, we observed that EVs derived from adult worms but not muscle larvae suppressed the expression of mucin-related genes from colon cell line and the organoids of small intestine. The mucin production in the intestine is thought to be enhanced by type 2 cytokines such as IL-13 when the helminth parasites are there. We found that AW-EVs suppressed steady-state expressions of mucin-related genes from the intestinal organoids. On the other hand, it is still unknown whether those EVs also suppress the enhancement of mucin-related gene expressions because we failed to induce such enhancement by exogenously adding IL-13 to the organoids. Moreover, it was difficult for us to evaluate the effect of EVs on mucin production in vivo, so that again it is still unknown whether the EVs act to suppress the expulsion of adult worms of *T. spiralis* in the infection. Further investigation is needed to address these issues.

Nurse cell formation and encapsulation by nurse cells are one of characteristics in *T. spiralis* infection. Although this phenomenon is triggered by the entrance of newborn larvae into the muscle cells and the following destruction of the muscle cells, its process is thought a combined reaction of host repairment in muscle and factors from the larvae inside of the muscle cells. However, the whole picture and mechanisms of nurse cell formation and encapsulation remain unclear. In this study, we showed that injection of ML-EVs but not AW-EVs into the muscle of uninfected mice induced the expression of genes related to myoblast differentiation. Moreover, our proteomics analysis revealed that muscle larvae-derived EVs contained proteins involved in the angiogenesis and VEGF signaling pathways, which are seen in the encapsulation of larvae in the muscles. From these findings, we think that the nurse-cell formation and encapsulation in the muscle of mice infected with *T. spiralis* might be regulated at least in part by EVs from larvae inside of muscles. There are several subspecies of *Trichinella* such as *T. pseudospiralis*, which induce incomplete encapsulation. The comparison in proteins of ML-EVs derived from between encapsulated *Trichinella*, such as *T. spiralis*, and unencapsulated *Trichinella*, such as *T. pseudospiralis*, will give us useful information to make clear the mechanisms underlying nurse cell formation and encapsulation.

Both AW-EVs and ML-EVs contains many miRNAs, such as tsp-miR-1, tsp-miR-100-5p, tsp-miR-125b-5p, tsp-miR-57-5p, and tsp-miR-72-5p, that are conserved among EVs secreted from other helminths ^33–36^. Among them, miR-1 is highly conserved among helminths and mammals. However, the function of helminth-derived miR-1 is limitedly reported. One study demonstrated that Schistosoma japonicum derived miR-1 causes hepatic fibrogenesis in mice by targeting Sfrp1gene ^37^. Our study elucidated that tsp-miR-1 could regulate the expression of murine *Il6* mRNA in murine RAW 264.7 cells. On the one hand, miR-36 family members are helminth-specific miRNAs and play critical roles in the early development of *Caenorhabditis elegans*. They also express in several helminths and are contained in helminth-derived EVs ^33,34,36^. Thus, the function of miR-36 in helminths might associate with their developmental processes seen in *C. elegans*. However, the function of miR-36 in the secreted EVs outside of the worm body is unknown. Surprisingly, we found in this study that tsp-miR-36-5p was very highly expressed in the adult worms with about 57-fold greater copy number/10^3^ *Gapdh* than that in muscles larvae, but this miRNA was only detected in ML-EVs (ML-EV specific miRNAs) with the highest abundance (Supplementary Table 3). It is very interesting to know why muscle larvae secrete tsp-miR-36-5p via EVs but not adult worms. It is possible that tsp-miR-36-5p might be secreted via EVs to modulate the host gene expressions in an unidentified process related to the muscle stage of infection. In general, the action of miRNA is quite dependent on the nucleic acid sequences. Even though we could show here that tsp-miR-1 suppressed the expression of murine *Il6* in the murine macrophage cell line, we did not find the target sequence of tsp-miR-1 in the homologous *Il6* gene in other organisms by searching in the database. According to this, miRNA in EVs might occasionally exert its function to other organisms but not always. However, it is more likely that proteins might exert same functions to other organisms compared to miRNA. Since protein-protein interactions can allow several tiny differences in the protein sequences over the organisms, it might be suitable for *T. spiralis* to use proteins in EVs to modulate host responses for their parasitic strategy.

We found that *T. spiralis* released EVs with stage-specific functions based on their contents such as proteins and miRNAs. From these findings, we speculated that *T. spiralis* might regulate the secretion of EVs in a stage-specific manner. We observed the difference in the biomolecules packed in the EVs between the two different infectious forms of *T. spiralis*, whereas we did not see any large difference in the expression of stage-specifically packed proteins and miRNA in the parasite bodies between of the two stages. We used whole bodies of parasites for the analysis of the expression of mRNAs and miRNAs. Although we did not identify the cells secreting EVs outside of the parasites, EVs should be secreted from the parasite organs including glands, esophagus, ventricle, intestine, and outer surfaces like excretory-secretory (ES) products. It is possible that the cells in those organs might stage-selectively express and pack the biomolecules in EVs. Higher resolution analysis at a single cell level is needed to be performed in future investigations.

In the present study, we comprehensively analyzed the extracellular vesicles derived from different infectious stages of *T. spiralis* and found that EVs derived from different stages of worms exert the common in both EVs and each EV-specific functions for their parasitism. These functions might be dependent on the different contents of biomolecules such as proteins and microRNAs. Although the whole functions of secreted EVs from *T. spiralis* are not clear, we believe that our findings are able to shed light on the progress in understanding trichinellosis.

## Methods

### Mice

Femal Slc:ddY mice (10 weeks old), male C57BL/6 mice (8 weeks old), and female BALB/c (6 weeks old) were purchased from Japan SLC (Hamamatsu, Japan). Femal Slc:ddY mice were used as the host for harvesting adult worms or muscle larvae for in vitro culturing to collect secretory products for EVs isolation. Male C57BL/6 mice were used as the host for maintaining *T. spiralis* life cycle for all experiments in our laboratory, and for in vivo effect of EVs on the myoblast differentiation. All mice were maintained at the Animal Resources Center of Gifu University Graduate School of Medicine under pathogen-free conditions. In case of mice used for generation of intestinal organoids, male C57BL/6 mice were purchased from CLEA Japan (Shizuoka, Japan). All mice care and experimental procedures were approved by the Committee for Animal Research and Welfare of Gifu University and were in full compliance with the institutional guidelines of Gifu University (No. 17179).

### *T. spiralis* stain and maintenance

Two hundred larvae/mouse *of T. spiralis* strain, ISS413, are infected to male C57BL/6 mice annually for maintaining their life cycle for use in our laboratory.

### Maintenance of cell lines

Cell lines including RAW 264.7 cells, HT-29 cells, and C2C12 cells were cultured in a 100 mm cell culture dish (TrueLine, USA) with growth media containing Dulbecco’s Modified Eagle’s medium (DMEM) (FUJIFILM Wako Pure Chemical corporation, Japan) supplemented with 10% fetal bovine serum (FBS) (biowest, Nuaille, France) and 1% penicillin/streptomycin (Gibgo) and maintained in a humidified atmosphere of 5% CO_2_ and 37°C. The cells were regularly passaged once high cell density was reached.

### Generation and maintenance of intestinal organoids

The small intestine of the mouse was dissected, washed in cold PBS, and divided into three parts (proximal, medial, and distal). The proximal part (~12 cm) including the duodenum and upper part of the jejunum was used for crypt isolation and organoid culture. Crypts were isolated as described previously ^38–40^. The isolated crypts were pelleted by centrifugation and washed with ice-cold basal culture medium (Advanced DMEM/F12; Thermo Fisher Scientific, Inc., Waltham, MA) supplemented with 2 mM glutamate, 10 mM HEPES, 100 U/ml penicillin, 100 μg/ml streptomycin. After centrifugation, the crypt pellet was resuspended in 200 µl of ice-cold basal culture medium, and the number of crypts was counted under an inverted microscope. A total of 500 to 1000 crypts/well were mixed with growth factor reduced Matrigel (Corning, Corning, NY) and plated in 48-well plates. Plates were incubated for 15 min at 37°C to allow the Matrigel to polymerize and then overlaid with 300 µl of growth medium (basal medium supplemented with 12.5 ng/ml mouse recombinant epithelial growth factor (PeproTech, Rocky Hill, NJ), 25 ng/ml mouse recombinant Noggin (PeproTech), 20% R-spondin1 (PeproTech) conditioned medium, and 5% fetal bovine serum) and incubated at 37°C, 5% CO_2_ for 7 days.

### Isolation of EVs from secretory products of *T. spiralis*

In order to collect secretory products from two stages of *T. spiralis,* muscle larvae and adult worms, the infective-stage larvae (muscle larvae) were infected to Slc:ddY mice by oral gavage. Adult worms were collected from the intestine at day 5 post-infection and muscle larvae were collected from striated muscle at day 30 post-infection. Parasites were cultured in serum-free media containing antibiotics for 2 days.^41^The culture media containing secretory products from parasites were harvested. Parasites and cell debris were removed by series centrifugation at 1000 × g, 5000 × g, and 10,000 × g. The supernatant was filtered through a 1-µm filter before purification of EVs by ultracentrifugation at 100,000 × g for 90 minutes. EVs pellet was washed with PBS by ultracentrifugation at 100,000 × g for 90 minutes. Finally, the EVs pellet was collected by re-suspension with PBS for functional analysis in cell lines, intestinal organoids, and mice. In the case of collection EVs for mass spectrometry and EV-Cell fusion analysis, RIPA lysis buffer or CFSE was added directly to the EVs pellet, respectively, after washing by PBS. EVs isolation protocol was modified from the previously report ^7^. EVs quantification was performed using Qubit™ Protein Assay Kit (Thermo Fisher Scientific,Waltham, Massachusetts, USA), then the EVs concentrations was read by Qubit®2.0 Fluorometer (Thermo Fisher Scientific, Waltham, Massachusetts, USA) ^42^. The EVs were prepared freshly prior to perform each experiment.

### Electron microscopy

Freshly prepared AW-EVs and ML-EVs suspension were dropped onto a carbon mesh for electron microscopic observation, then allow drying at room temperature for 5 minutes. Excess water was removed with filter paper and the carbon mesh was immerse in 1% phosphotungstic acid for 10 seconds. Then excess water was removed with filter paper and dried out at room temperature for 20 minutes. Then the EVs were visualized by electron microscopy.

### Analysis of EV-Cell fusion

To confirm whether the existing EV-like particles identified by TEM are membrane-surrounded particles and capable to fuse with the plasma membranes. ML-EVs were obtained by ultracentrifugation after removing supernatant EVs pellet were labeled with 45 µM of CSFE (Thermo Fisher Scientific, Waltham, Massachusetts, USA) at 4 °C for 30 minutes, then CFSE-labeled EVs were washed in PBS by ultracentrifugation to remove excessive CFSE. CFSE-labeled EVs were added to HT-29 cells and incubated at 4 °C to let EV fuse with the plasma membranes, for 30 minutes then EV-Cell fusion was determined by FACS analysis ^43^.

### Functional analysis of EVs on cell lines

#### RAW 264.7 cells

To test the effect of EVs on macrophage activation, RAW 264.7 cells were seeded onto tissue culture plate 96 wells total of 5,000 cells/well. The cells were grown for 24 hours, AW-EVs or ML-EVs at the concentration of 50, 25, and 12.5 µg/ml were added and continued culture for another 24 hours then the cells were stimulated by 100 ng/ml LPS for 6 hours. A culture medium was subsequently collected to examine secreted IL-6 by ELISA. In order to measure *Il6* mRNA expression in RAW 264.7 cells, the cells were seeded onto tissue culture plate 48 wells total of 12,000 cells/well culture for 24 hours, then 12.5 µg/ml of AW-EVs and ML-EVs were added. The cells were continued cultured and stimulated by LPS same as the culture for IL-6 secretion. Then cells were harvested for total RNA isolation and RT-qPCR. For detecting the expression of CD80 by fluorescence-activated cell sorting (FACS) analysis, cells were cultured the same as for measuring *Il6* expression but different in EV concentrations (for CD80 detection 50 and 25 were used). Then cells were harvested for FACS analysis.

#### HT-29 cells

In order to test the specific function of AW-EVs, HT-29 cells total of 20,000 cells/well were seeded onto tissue cell culture plate 24 wells and cultured for 24 hours, then 12.5 µg/ml AW-EVs, 12.5 µg/ml ML-EVs or PBS were added and continued cultured for 24 hours. Total RNA was isolated for RT-qPCR to measure the mRNA expression of mucins genes.

#### C2C12 cells

To test the specific function of ML-EVs on myoblast differentiation, C2C12 cells (10,000 cells/well) were grown in normal growth media in a 48-well plate for 24 hours or until cells reached 80% confluency. The cells were induced to differentiation by culture in differentiation media (DMEM supplemented with 0.5% FBS) in the present of 12.5 µg/ml AW-EVs, 12.5 µg/ml ML-EVs or PBS. At day 2, 4 and 7 after differentiation, cells were harvested for measurement of expression of genes related to differentiation of myoblast, *Myog* and *Myh3* ^20,44,45^.

### Effect of EVs on intestinal organoids

After 7 days of culture, organoids were passaged by dissolving Matrigel domes in ice-cold PBS, transferred into 15-ml tubes, and centrifuged at 200 × g for 2 minutes The supernatant was discarded, the pellet was resuspended in 3 ml of ice-cold Advanced DMEM/F12, and organoids were disrupted by strong pipetting 20 times with a 1000-µl pipette tip. After centrifugation at 200 × g for 2 minutes and removing the supernatant, 1 ml of growth medium was added to the pellet and then crypt number was counted. The 24-well plates were coated with 4% Matrigel dissolved in ice-cold PBS and incubated at 37°C for 1 hour. The plates were washed three times with PBS at room temperature immediately before use. The crypt suspension was diluted with growth medium and about 1500 crypts/0.5 ml were dispensed to each well. Crypts were incubated at 37°C, 5% CO2 for 2 days, then 6 µg/ml of AW-EVs or ML-EVs in growth media were added to replace old growth media. Intestinal organoids were continued culture in the presence of EVs for 24 hours. Total RNAs were collected to examine the influence of both EVs on mucin-related genes, *Tff3* and *Clca3*.

### *In vivo* effect of EVs on myoblast differentiation

To investigate the specific function of ML-EVs on the myoblast differentiation in vivo, a Total of 7.5 µg of AW-EVs or ML-EVs in 50 µl PBS were injected to the tibialis anterior muscle of mice. Two days after injection, 2 g of EV-injected muscle was collected. The collected muscle was homogenized by OMNI TIP™ HOMOGENIZING KITS with stainless steel generator probe (Omni International, USA), at the speed of 25,000 rpm for 40 seconds in 500 µl of RNAiso plus. Expression of *Myog*, *Mrf4,* and *Myh3* were measured by RT-qPCR.

### RNA isolation and quantitative reverse transcription polymerase chain reaction (RT-qPCR)

To determine the mRNA and miRNA expression of each experiment. Total RNA was isolated from a treated cell line, muscle tissue, adult worm, muscle larvae, and small intestinal organoids by Total RNA extraction reagent RNAiso Plus (TaKaRa Bio, Shiga, Japan) or NucleoSpin^®^RNA (MACHEREY-NAGEL, Duren, Germany) according to manufacturer’s instruction. Reverse transcription was performed by ReverTra Ace™ qPCR RT Master Mix with gDNA remover (TOYOBO,Osaka,Japan) or ReverTra Ace^®^ qPCR RT Master Mix (TOYOBO,Osaka,Japan) according to manufacturer’s instruction. In case of miRNAs, reverse transcription was performed as previously described ^46^. In brief, RNA was polyadenylated with Poly(A) Polymerase Tailing Kit (BIOSEARCH TECHNOLOGIES). Then the polyadenylated product was reversely transcribed into cDNA using a poly(T) adapter (3’CAGTGCAGGGTCCGAGGTCAGAGCCACCTGGGCAATTTTTTTTTTTVN-5’) with a ReverTra Ace qPCR RT Kit (TOYOBO, Japan). Real-time PCR was performed using THUNDERBIRD Next SYRB qPCR Mix (TOYOBO) on a real-time PCR system (Thermal Cycler Dice® Real Time System, Takara).

For miRNAs expression differentiation and level in adult worms and muscle larvae, absolute quantitative real time PCR was performed as our previously described.^47^ Twenty microliters of the reaction solution consisted of 2 μl cDNA (5-time dilution), 0.8 μl of 5 M forward primer (miRNA specific primer), 0.8 μl of 5 uM reverse primer (universal primer, 3’-CAGTGCAGGGTCCGAGGT-5’), and 10 μl of SYBR reagent. PCR amplification was performed with a denaturation step at 95 °C for 5 s and annealing and extension at 60 °C for 30 for 45 cycles.

Specific external controls were constructed for all target miRNAs. The PCR fragment of each miRNA was cloned into a pT7Blue T-Vector (Novagen, Inc, Madison, WI, USA). The recombinant plasmids were introduced into competent cells of *Escherichia coli* JM 109. The plasmid DNA was isolated from *E. coli* using a PureYield™ Plasmid Miniprep System (Prometa, Madison, USA).

Copy number of miRNAs were calculated based on molecular weight and concentration of the recombinant plasmid. Tenfold serial dilutions (10^1^ to 10^7^ copies/2 μl) of the plasmids were used to generate standard curve for each miRNA. Differences in the amounts of cDNA from different samples were normalized by housekeeping gene glyceraldehyde 3-phosphate dehydrogenase (*Gapdh*), and expression levels were represented as the copy numbers of target miRNA/10^3^ *Gapdh* copies. Log ten bases was applied to copy number of target miRNA/10^3^ *Gapdh* copies to reduce variation between samples. The data of miRNAs expression in adult worms and muscle larvae are shown as log (copy numbers/10^3^ *Gapdh* copies.

Relative expression of other genes was calculated by the Delta-Delta Ct method (2^−ΔΔCT^ method). Briefly, target mRNAs of each sample were calculated by normalizing to their housekeeping gene mRNAs (mouse and murine cell lines used *Gapdh* as housekeeping. For the human cell line, *GUS* was used). The normalized target mRNAs of each sample were then relative to the normalized target mRNAs of control samples. The control sample of each experiment was explained in each figure legend. The relative expressions were presented as fold gene expressions. In the case of the expression of mRNA of EV proteins in adult worms and muscle larvae, the mRNA expression level was calculated by 2^-(ΔCT)^. Where ΔCT = Ct of target mRNA-Ct of *Gapdh* mRNA. The 2^-(ΔCT)^ of each target mRNA was multiplied by 10000 to make all 2^-(ΔCT)^ values greater than 1. Log ten bases was applied to 2^-(ΔCT)^ x 1000 to reduce variation between samples. Thus, the level of mRNA of EV proteins in adult worms or muscle larvae are the result of log_10_(2^-(ΔCT)^ × 1000). In Figs. 4b-d, log_10_(2^-(ΔCT)^ ×1000) were denoted as log(2^-(ΔCT)^). Primer sequences were listed in Supplementary Tables 4-6.

### Enzyme-linked immunosorbent assay (ELISA)

To measure the amount of IL-6 secreted by RAW 264.7 cells, after co-cultured with EV and stimulated by LPS, cell culture supernatant was collected. The high protein-binding capacity 96 well ELISA plate (Nunc MaxiSorp® flat-bottom, Invitrogen) was coated with ELISA capture antibody, purified anti-mouse interleukin-6 (IL-6) (MP5-20F3) with 1:1000 dilution, after incubation at 4 °C overnight, the plates were washed 2 time with PBS containing 0.05% Tween-20 then blocked with 1% bovine serum albumin (BSA) in PBS (blocking buffer) for 1 hours at room temperature. After blocking, the plates were washed and the culture supernatant with dilution 1:1 was added to the plates, after 2 hours of incubation at room temperature, plates were washed, and the detection antibody, Biotin anti-mouse IL-6 (MP5-32c11) with the dilution 1:500 were added to the plates and incubated at room temperature for 90 minutes. To visualize the detection antibodies, the plates were incubated with horseradish peroxidase (HRP) for 45 minutes at room temperature, after that 3,3’,5,5’-tetramethylbenzidine (TMB) (Sigma-Aldrich) as substrate of HRP were add to the remaining HRP after washed-plates. The reaction was stopped by adding of 2 M sulfuric acid. The plate reader (SUNRISE RAINBOW) was used for measure OD at 450 nm. The concentrations of IL-6 were calculated by PLATEmanager V5 ®software (FUJIFILM, Japan)

### FACS analysis

To evaluate expression of CD80 in RAW 264.7 cells, after treated with AW-EVs or ML-EVs and stimulated by LPS. Cells were harvested by trypsinization with 2.5% trypsin, the reaction was stopped by FBS then cell were washed 2 time with PBS-based staining buffer containing 2% fetal bovine serum (FBS) and 0.01% NaN3 (Sigma-Aldrich), blocked Fc IgG receptors by pre-incubated with CD16/32(93) at 4 °C for 15 minutes, then washed and stained with specific monoclonal antibody to cell surface antigen including CD11b-PE(M1/70), CD80-APC (16-10A1) at 4 °C for 20 minutes, then cells were washed and FACS analysis was performed by CytoFLEX S Flow Cytometer (Beckman Coulter Life Sciences). The data was analyzed by FlowJo (Tree. Star, OR) software. The concentrations of antibodies used in this study followed the manufacturer’s instruction.

### Mass spectrometry and bioinformatic analysis

Proteomic analysis of AW-EVs and ML-EVs were performed using LC-MS/MS as described in previous study ^48^. Briefly, total 500 of RIPA lysis buffer were added directly to AW-EVs or ML-EVs pellets after isolation by ultracentrifugation. The protein samples were prepared before trypsin digestion by precipitated with methanol-chloroform, dissolved into RapiGest™ (Water corporation, UK), reduced with 10mM dithiothreitol (DTT), followed by alkylation with iodoacetamide. After trypsin digestion, the peptides were purified with C18 tip (GL-Science, Tokyo, Japan). Then subjected for LC-MS/MS analysis.

Proteins were identified by MASCOT Server (Matrix Science, London, UK; version 2.7.0). Mascot was set up to search against the Uniprot-Trichinella-spiralis_20210607 database. Scaffold software version 5.1.2 (Proteome Software, Portland, OR, USA) were used for calculating the quantitative value of each protein.^49^

Three categories of EV proteins, including common EV proteins, AW-EVs specific proteins, and ML-EVs specific proteins were classified based on the existing of proteins in AW-EVs or ML-EVs. common EV proteins are proteins that identified in both AW-EVs and ML-EVs at least 1 biological replicate, AW-EV specific proteins are proteins that only identified in AW-EVs at least one biological replicate but not in ML-EVs, ML-EV specific proteins are proteins that only identified in ML-EVs at least one biological replicate but not in AW-EVs.

Functional analysis of protein based on Gene ontology analysis was analyzed by The PANTHER classification system (www.pantherdb.org) as previous described ^50^. Briefly, Uniprot accession number of proteins were prepared and uploaded, *T. spiralis* was selected as organism for id list, functional classification was analyzed in 2 aspects including, Biological process and Panther pathway.

### Whole miRNA sequencing and bioinformatic analysis

In order to isolate small RNAs from EVs. AW-EVs or ML-EVs were isolated from parasite culture media using Total Exosome Isolation (from cell culture media) (Invitrogen, USA). Small RNA was isolated from AW-EVs or ML-EVs pellets using NucleoSpin^®^ miRNA (MACHEREY-NAGEL, Duren, Germany). The small RNA library was prepared using NEBNext Small RNA Library Prep Set for Illumina (New England Biolaps, Ipswich, MA). Adapter was trimmed from reads by cutadapt. The sequencing reads were mapped to *T. spiralis* genome (GCF_000181795.1_Trichinella_spiralis-3.7.1) using mapper.pl. Then the reads were analyzed for frequency and normalized count. miRDeep2 (version 2.0.1.3) was used for predicted miRNAs and quantified read counts. miRNAs were identified by mapped all mature miRNAs sequences against all organism in miRBase database.

miRNAs identified in AW-EVs and ML-EVs were classified into 3 categories including common EV miRNAs, AW-EV specific miRNAs and ML-EV specific miRNAs. The criteria for classification are same as criteria of EV proteins classification.

Target miRNAs were predicted by TargetScan software and miRSystem database.

### miRNAs synthesis and transfection

Forward and reward sequences of tsp-miR-1, tsp-miR-5437, tsp-miR-107, tsp-36-3p, tsp-miR-181b-5p, and miRNA negative control were synthesized by Hokkaido System Science (Sapporo, Japan). List of miRNAs sequences are provided in supplementary Table 7. Each miRNA was hybridized into duplex by combine forward and reverse sequences in annealing buffer containing 30mM HEPES-KOH pH 7.4, 100 mKCl, 2mM MgCl2 and 50mM NH4Ac. The solution was incubated at 90 °C for 1 minute and slowly cooled down to room temperature over a period of 45 minutes. Twenty-four hours before transfection, RAW 246.7, HT-29 and C2C12 cells were grown in normal growth media, then 7.5 nM of each synthetic miRNA were transfected into cells by Lipofectamine® RNAiMAX Reagent (Invitrogen, USA) as manufacturer’s description for 24 hours. In case of HT-29 cells, after transfection, total RNA was collected to examine *MUC5AC* mRNA expression. For RAW 264.7 cells, they were stimulated by 100 ng/ml of LPS for 6 hours. Then total RNA and supernatant were harvested to examine expression of *Il6* mRNA and protein expression respectively. For C2C12 cells, they were culturedin differentiation mediafor 7 days. At 2, 4, and 7 days after differentiation, total RNA were isolated to examine expression of *Myog* and *Myh3* mRNAs.

## Statistical analysis

The statistical detail of experiments was explained in figure legends. The data were analyzed using GraphPad Prism Version 9 for windows, (GraphPad Software, La Jolla, California, USA). For multiple group comparisons, the data were test for normal distribution by Shapiro-Wilk test. The data that passed (*P*≥0.05) or not passed (*P*<0.5) normality test was then analyzed using one-way ANOVA test or one-way ANOVA with Kruskall-Wallis test respectively. Statistically significant was assigned as ns, not significant. *p*<0.05. **, *p*<0.01. ***, *p*<0.001. ****, *p*<0.0001.

## Acknowledgement

We thank Y. Kouyama for her secretarial assistance. This work was supported by JSPS KAKENHI Grant Number 20H03478.

## Author contribution

S.K., Z.W., and Y.M. designed the study. S.K. and Z.W. carried out the biological analyses. R. H-Y. and T. Y. provided intestinal organoids. T. O. performed proteomics and whole microRNA sequencing analyses. C. T., H.O., and S. O. performed electron microscopical analysis. S.K. and Y.M. wrote the manuscript. All contributors approved and edited the manuscript.

