## Supplemental Figures 1, 2. Supplemental Tables 1-7. for "Trichinella spiralis secretes stage-specific extracellular vesicles for modulating host microenvironment for parasitism"

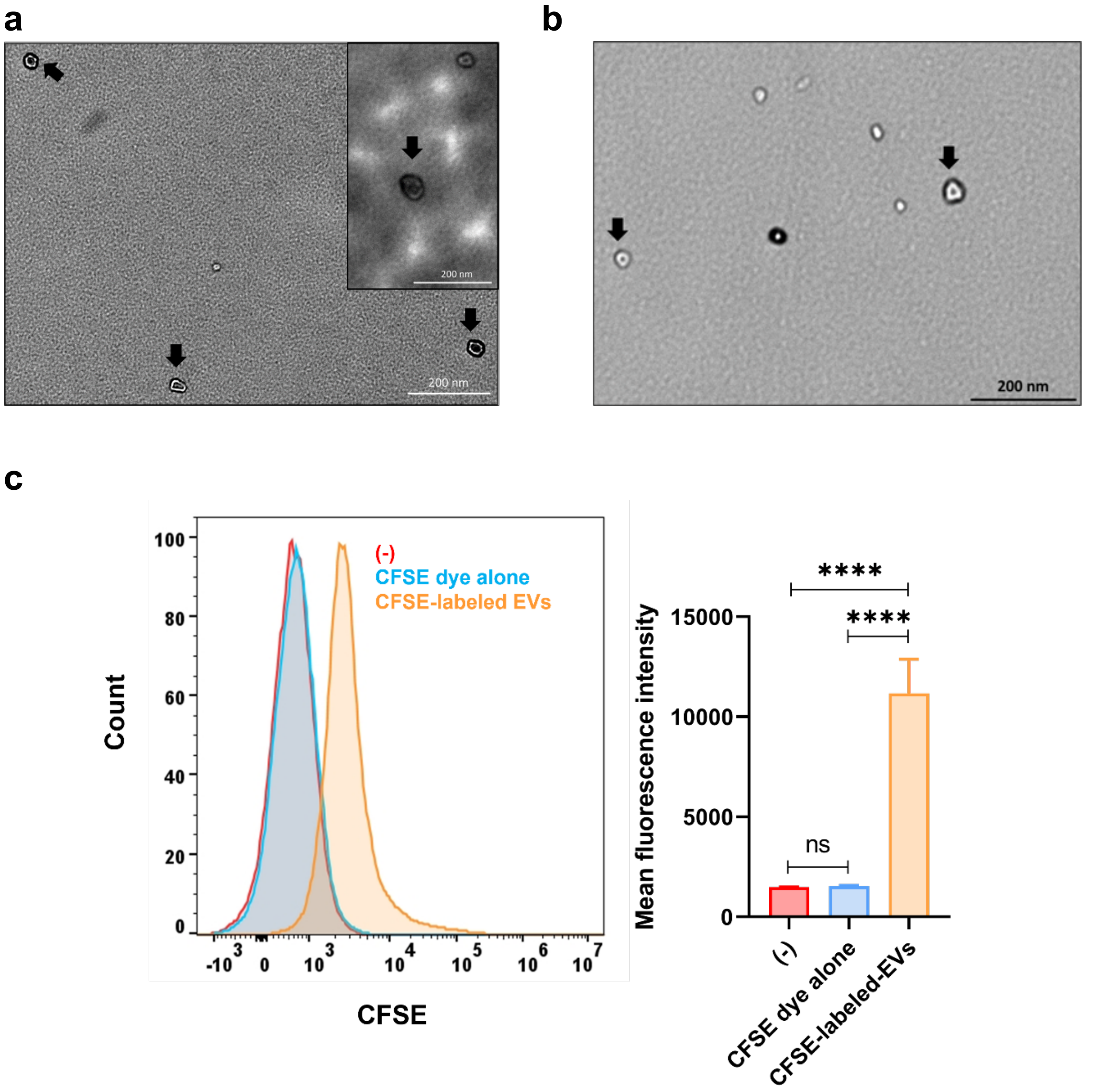


**Supplementary Fig. 1 Characterization of extracellular vesicles secreted from *T. spiralis* adult worms and muscle larvae by electron microscopy and flow cytometry.**

**(a, b)** Ultrastructure of AW-EVs **(a)** and ML-EVs **(b)** were visualized by electron microscopy. The black arrows indicate EVs-like structure.

**(c)** Mean fluorescence intensity (MFI) of CFSE in HT-29 cells were measured by flow cytometry. red, blue, and orange histogram indicate MFI of CFSE in HT-29 cells that were incubated in growth media alone (-), co-incubated with CFSE dye alone and co-incubated with ML-EVs that were labeled by CFSE (CFSE-labeled EVs) respectively at 4ºC. The data are representative of two independent experiments.

Graph presents mean values ± SD (n=3) (**b**). ns, not significant ****, $p<0.0001$. The statistically significant were calculated using one-way ANOVA (**b**).

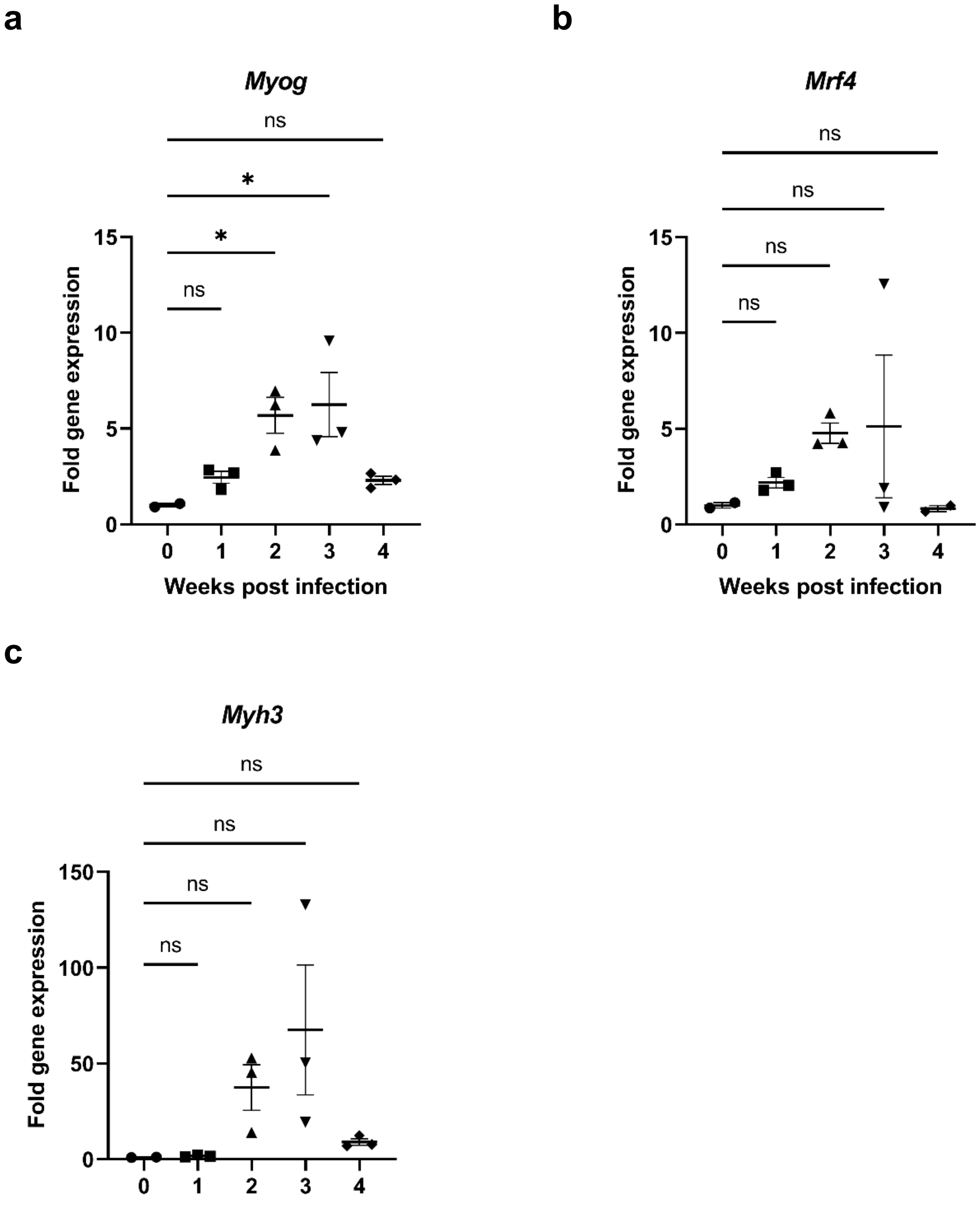


**Supplementary Fig. 2 Expression of genes related to differentiation of myoblast in *T. spiralis* infected mice**

Mice were infected with *T. spiralis* muscle larvae (200 muscle larvae/mouse). At 1-, 2-, 3- and 4-weeks post infection, tibialis anterior muscle was collected to examine expression of *Myog* (**a**), *Mrf4* (**b**) and *Myh3* (**c**). The normalized target mRNAs of each sample are relative to normalized target mRNAs of un-infected mice ((-)). The relative expressions are presented as fold gene expression.

Each dot represents an individual mouse. Error bars denote mean±SEM.

ns, not significant. *, $p<0.05$. The statistically significant were calculated using one-way ANOVA test (**a** and **c**) or one-way ANOVA with Kruskall-Wallis test (**b**).

### Supplementary Table 1 Representatives of common EV proteins and top ten most abundant AW-EV specific, and ML-EV specific proteins

| Categories of EV proteins | Protein name | Accession Number | Mean of Quantitative Value (SD) | |
|---|---|---|---|---|
| | | | AW-EVs | ML-EVs |
| Common | Putative serine protease | B0F9U0 | 30.82 (3.89) | 23.53 (6.06) |
| | Cystatin-like protein | B0FJW6 | 63.52(14.41) | 49.86 (9.55) |
| | ADP-ribose pyrophosphatase, mitochondrial | A0A0V1ASG0 | 66.81(19.39) | 53.44(12.30) |
| | Putative nudix hydrolase 6 | A0A0V1B1Q3 | 40.13(1.40) | 33.94 (3.47) |
| | ADP-ribose pyrophosphatase, mitochondrial | A0A0V1ARQ9 | 41.86(15.87) | 36.70 (10.69) |
| | Uncharacterized protein ad48h-Ts3 | F0V432 | 42.59(1.60) | 40.21 (11.92) |
| | Mannose-6-phosphate isomerase | A0A0V1BCS6 | 19.61(1.84) | 18.60 (5.75) |
| | Chymotrypsin B | A0A0V1B629 | 10.89(3.57) | 10.77 (4.22) |
| | L-cystatin | A0A0V1B7I5 | 15.08(2.98) | 18.86 (7.57) |
| | Tissue-type plasminogen activator | A0A0V1AVL6 | 25.89(1.36) | 32.54 (8.29) |
| AW-EVs | Deoxyribonuclease-2-alpha | A0A0V1B6X3 | 44.85(7.6) | 0.00 |
| | ZP domain-containing protein | A0A0V1BSB5 | 43.81(10.03) | 0.00 |
| | Peptidase S1 domain-containing protein | A0A0V1BXM4 | 26.52(3.86) | 0.00 |
| | Adult-specific DNase II-3 | Q32R73 | 26.30(2.94) | 0.00 |
| | Adult-specific DNase II-4 | Q32R72 | 26.30(0.58) | 0.00 |
| | Adult-specific DNase II-12 | A0A165G716 | 23.82(4.95) | 0.00 |
| | Newborn larvae-specific serine protease SS2-1 | Q6RUJ3 | 23.12(3.39) | 0.00 |
| | Major sperm protein | A0A0V1BX53 | 22.51(6.43) | 0.00 |
| | Newborn larvae-specific DNase II-4 | Q32VV2 | 18.49(6.89) | 0.00 |
| | Newborn larvae-specific DNase II-3 | Q32VV3 | 18.35(4.82) | 0.00 |
| ML-EVs | Cuticlin-1 | A0A0V1BL36 | 0.00 | 34.27 (19.35) |
| | Cuticlin-1 | A0A0V1BD95 | 0.00 | 21.31 (16.75) |
| | Transmembrane matrix receptor MUP-4 | A0A0V1BTK0 | 0.00 | 13.25 (8.86) |
| | BPI2 domain-containing protein | A0A0V1B5A1 | 0.00 | 12.87 (3.12) |
| | Uncharacterized protein | A0A0V1BTN2 | 0.00 | 9.77 (2.39) |
| | Metalloendopeptidase | A0A0V1C237 | 0.00 | 9.54 (1.26) |
| | Uncharacterized protein | A0A0V1AZ10 | 0.00 | 9.10 (9.12) |
| | Uncharacterized protein | E5SAW8 | 0.00 | 8.95 (5.37) |
| | Uncharacterized protein | E5SYG4 | 0.00 | 8.04 (3.86) |
| | Alpha-1,4 glucan phosphorylase | A0A0V1AU99 | 0.00 | 7.96 (5.89) |

**Supplementary Table 2. List of EV protein markers and proteins found in EVs derived from other helminths sharing with *T. spiralis* AW-EVs and ML-EVs.**

| Protein names | Accession number | References |
|---|---|---|
| 14-3-3 protein | C9DTY3 | [1–6] |
| Heat shock protein 70[a] | Q961X6 | [1,4–9] |
| Actin-5C | E5SLU1 | [3–5,7] |
| Cathepsin L | A0A0V1BTW2 | [3] |
| Glyceraldehyde-3-phosphate dehydrogenase 1[a] | A0A0V1C2A5 | [1] |
| L-lactate dehydrogenase | A0A0V1AYM5 | [1,6] |
| Fructose-bisphosphate aldolase | A0A0V1AX39 | [1,3,6,8] |
| Elongation factor 1-alpha | A0A0V1BIW5 | [1,7,8] |

List of proteins identified in AW-EVs and ML-EVs that common with EV proteins markers in ExoCarta database or EV proteins from other helminths.

[a] Proteins that are classified as EV proteins markers in ExoCarta database which is an open- source website containing information of proteins and RNAs identified in exosomes originated from several cell types.

**Supplementary Table 3. Representatives of known common EV miRNAs and top ten most abundant AW-EV specific and ML-EV specific miRNAs.**

| Categories of EV miRNAs | miRNA name | Mature sequence | Normalized count | |
|---|---|---|---|---|
| | | | AD-EVs | ML-EVs |
| Common | tsp-miR-100-5p | aacccguagauccgaacuugugu | 2024640.6 | 465794.27 |
| | tsp-miR6235-3p | aacccguauuuuccucuuagucu | 82889.113 | 36585.677 |
| | tsp-miR-124-5p | cguguucacugucugucuugau | 5487.9893 | 5933.5702 |
| | tsp-miR-56 | uacccguaucucuucuugguu | 3344.7273 | 6150.8474 |
| | tsp-miR-53-5p | uacccguaucuuucuuucaguau | 10632.418 | 20141.697 |
| | tsp-miR-72-5p | aggcaagauguuggcauagcuga | 11199.693 | 39915.926 |
| | tsp-miR-263a-5p | cuuggcacuguaagaauucaca | 29465.359 | 125500.12 |
| | tsp-miR-2238l-3p | uggacggcgaauuaguggaag | 1780.7252 | 7627.7316 |
| | tsp-miR-1 | uggaauguaaagaaguauguag | 22426.351 | 118416.08 |
| | tsp-miR-228-5p | aauggcacuggaugaauucacgg | 53853.205 | 355073.01 |
| AW-EVs | tsp-miR-5437 | gagauaaaguggaacggucauu | 31221.116 | 0 |
| | tsp-miR-3352 | ucuucuucuguucuauaucguguuu | 1429.1743 | 0 |
| | tsp-miR-35a-3p | uggaaaaauugagccaugguuau | 1006.714 | 0 |
| | tsp-miR-7883a-5p | uggcuguacaucguggacuuuac | 742.05206 | 0 |
| | tsp-miR-1298-3p | ucuuccguccgucgcccaguuguau | 528.32509 | 0 |
| | tsp-miR-497-3p | uuggcacugcagcucaccacgc | 521.33402 | 0 |
| | tsp-miR-2238i-3p | ugccgcacuguugaucuucaggu | 432.44757 | 0 |
| | tsp-miR-7794 | ucgugcuuuagaauccgguaguu | 387.50498 | 0 |
| | tsp-miR-2839-3p | cucuuccgcgcacuguuugauga | 291.62746 | 0 |
| | tsp-miR-382-3p | uucacggauagcacuuuugguagc | 284.63639 | 0 |
| ML-EVs | tsp-miR-36-3p | ucaccggguaauaauucacagc | 0 | 9531.1601 |
| | tsp-miR-3084b-3p | ugccaguucccuccugaacacu | 0 | 7361.3918 |
| | tsp-miR-4752 | ucgaucauccuugagaaccaga | 0 | 4279.4598 |
| | tsp-miR7510a | ugggacgcuaaagaucuugagga | 0 | 2854.642 |
| | tsp-miR-2004 | auguuccugugguugcgca | 0 | 1449.8498 |
| | tsp-miR7787-3p | ucuucggucuguacuugguagga | 0 | 1123.4333 |
| | tsp-miR-345-3p | aagacuagaggucaggagaaug | 0 | 860.09733 |
| | tsp-miR-993 | gaagcucguuucuacaggcaugu | 0 | 708.90444 |
| | tsp-miR-27a-5p | uuaccugucuugugaaugcgac | 0 | 315.4024 |
| | tsp-miR-9769-3p | uuuucuguucugugugcuugacu | 0 | 137.17501 |

**Supplementary Table 4. Primer sequences of mouse and human for RT-qPCR.**

| Gene | Species | Primer | Sequence |
|---|---|---|---|
| *Gapdh* | Mouse | Forward | CCCGTAGACAAAATGGTGAAGG |
| | | Reverse | GGCATTGTGGAAGGGCTCAT |
| *Il6* | Mouse | Forward | TTCCATCCAGTTGCCTTCTTG |
| | | Reverse | CATTTCCACGATTTCCCAGAG |
| *Myog* | Mouse | Forward | AACTACCTTCCTGTCCACCTTCA |
| | | Reverse | TCCCCAGTCCCTTTTCTTCC |
| *Mrf4* | Mouse | Forward | TAAGGAAGGAGGAGCAAACG |
| | | Reverse | CTGAAGAAATACTGTCCACGATG |
| *Myh3* | Mouse | Forward | CAGAGTGAGGAGGACAGGAAG |
| | | Reverse | CTGAACTTGGTGAGATGAGCA |
| *Clca3* | Mouse | Forward | ACGGGTGTCTGTGTTCATCC |
| | | Reverse | GCCAGGAATGGTGCTGTTG |
| *Tff3* | Mouse | Forward | CGGCAAATGTCAGAGTGGA |
| | | Reverse | TTGAAGCACCAGGGCACA |
| *GUS* | Human | Forward | ATGGAAGAAGTGGTGCGTAGG |
| | | Reverse | AAGGATTTGGTGTGAGCGA |
| *MUC1* | Human | Forward | GAATACCCTACCTACCACACTCACG |
| | | Reverse | TTACCTGCCGAAACCTCCTC |
| *MUC5AC* | Human | Forward | GCTGAGAGAAGACGGATGCTG |
| | | Reverse | GCAGTTTTGCTGACGGATGA |

### Supplementary Table 5. Primer sequences of mRNA of *T. spiralis* EV proteins

| Protein name | Accession number | Primer | Sequence |
|---|---|---|---|
| 14-3-3 protein zeta | C9DTY3 | Forward | GACCAACGAGGAACGCAAC |
| | | Reverse | CCTTCAGTTTTCTGCTCAATGC |
| ADP-ribose pyrophosphatase, mitochondrial | A0A0V1AVP2 | Forward | GGTTTCGTTCCTGATGCTC |
| | | Reverse | TGTGCCTGACTTGGTTTCTG |
| ADP-ribose pyrophosphatase | A0A0V1ARQ9 | Forward | CGCACCACCTGATTTTTCTG |
| | | Reverse | ATGCCCGTTTATGCTTCGTC |
| Chymotrypsin B | A0A0V1B629 | Forward | CAGTGAAATGGGAAGGCAAG |
| | | Reverse | CTCAGACAAGCAGAGCATACG |
| Collagen alpha-6 (VI) chain | E5SP50 | Forward | TTTACCCGACAGAATGTTTCC |
| | | Reverse | TCATCACTCACGCTTCCTG |
| Collagen alpha-6(VI) chain | A0A0V1BCC9 | Forward | TTTACCCGACAGAATGTTTCC |
| | | Reverse | TCATCACTCACGCTTCCTG |
| Cyclin-K | A0A0V1C002 | Forward | TGTGGTCAATCTGGGAAAAC |
| | | Reverse | AGAGGCTTGGGCTGGATAA |
| Cystatin-like protein | B0FJW6 | Forward | TCCACCAGACAAGTTCACCAG |
| | | Reverse | AAGGGATGAGACGGCAACC |
| Mothers against decapentaplegic homolog | A0A0V1BXV2 | Forward | GCTTTGGTCCGCCTATGTG |
| | | Reverse | TGCTTCGGGCAGTATCTTG |
| Cytohesin-3 | A0A0V1BK21 | Forward | TCCACCAGACAAGTTCACCAG |
| | | Reverse | AAGGGATGAGACGGCAACC |
| Gamma-aminobutyric acid receptor subunit pi | A0A0V1BAW3 | Forward | TCGTGTTGGCGGTTTGAATA |
| | | Reverse | GTGTTAGTTTGTCGGCTTTCCA |
| Glucose dehydrogenase | A0A0V1C2P7 | Forward | GCTGAAAACGAAAGCGAACTC |
| | | Reverse | TGGCAAACAACCACCAAGT |
| L-cystatin, partial | A0A0V1B7I5 | Forward | GTGTCAGCACAGGATGCACA |
| | | Reverse | AGGCTTCTCGTGGATGGTAAC |
| Low-density lipoprotein receptor-related protein | A0A0V1B0U4 | Forward | TTTTCGGTCTGGCGTTGAC |
| | | Reverse | GATGCGGCTTGGTGGATT |
| Mitochondrial-processing peptidase subunit alpha | A0A0V1BGF6 | Forward | GGGTTTGCCTAAGTTCTGTCC |
| | | Reverse | TCCCACTCCTCCAACAACAA |

### Supplementary Table 5. continued

| Protein name | Accession number | Primer | Sequence |
|---|---|---|---|
| Protein HIRA | A0A0V1ASJ6 | Forward | GCATTAGTTGGCATCAGTTCAG |
| | | Reverse | CCTGTCTGTTTGGCGTAAGG |
| Myoglobin | B0F9T0 | Forward | GTGGATTTTCGTTTTCATCACC |
| | | Reverse | TCCGCCGTTTTCGTTGTT |
| Neprilysin-2 | A0A0V1B9I8 | Forward | GGACGAGACAGGGTTATTGGTT |
| | | Reverse | ACATCTTTTTGGCGGATTCA |
| Papilin | A0A0V1BLP2 | Forward | GCGAAGGAAACGGCAATC |
| | | Reverse | CCAAACAAGGTCCAGCATCA |
| Plancitoxin-1, partial | A0A0V1BVT6 | Forward | ACCCTACGCCCTTACTATGCTG |
| | | Reverse | GGAATGGAAACTGTGGACGA |
| Putative serine protease | B0F9U0 | Forward | TACTGCCCGAAGCATTGA |
| | | Reverse | CTTTCTTTCCAACATCTACTCTGC |
| Putative nudix hydrolase | B0F9S5 | Forward | TCTGCCTGGTGAGGAATCG |
| | | Reverse | CAGCGTCAAAGAAAGCACTGTA |
| Sodium/potassium-transporting ATPase subunit beta-1 | A0A0V1BN13 | Forward | CCCCTGCTAAATCTCTCTTCCA |
| | | Reverse | CGCACCTCGTCTGTTCAAA |
| Putative tyrosinase-like protein tyr-3 | A0A0V1BH07 | Forward | CAGTTTTGGGCCTCAGTCG |
| | | Reverse | GGAAAGCATGTGTTGCAGGA |
| Snake venom 5'-nucleotidase | A0A0V1BJP9 | Forward | GCGTTCATTCCCTACAACTACG |
| | | Reverse | AATCATCTCCGCCTTTAGCC |
| Tissue-type plasminogen activator | A0A0V1AVL6 | Forward | TGTATCTGGCAAAAGGTGGA |
| | | Reverse | TGCAACGTGTAGTCGTCAGC |
| Uncharacterized protein T01_6155 | A0A0V1BX36 | Forward | CCACTACCGTGCGTTGATACA |
| | | Reverse | CCATTGCTGTCTACTTTGATTTTCC |
| Uncharacterized protein T01_7775 | A0A0V1BIH7 | Forward | CCCAAAGATTGCCGACCA |
| | | Reverse | GTTTTCCATCAGGCGAAGGT |
| Uncharacterized protein | E5RYD6 | Forward | GGATGCCAGAGCACAACGTA |
| | | Reverse | CCCCAATCACACACTGATGG |
| Venom allergen 5 | A0A0V1BWX2 | Forward | ACTGGTGAGGGTTCTGACAA |
| | | Reverse | GCACATTTTCCTTCTTTTCTCC |

### Supplementary Table 5. continued

| Protein name | Accession number | Primer | Sequence |
|---|---|---|---|
| ZP domain-containing protein | A0A0V1BSB5 | Forward | CCGATGGAACGAAACGAAA |
| | | Reverse | CCAACAGCGAACAGTGGATG |
| Adult-specific DNase II-12 | A0A165G716 | Forward | TTACACACCTGGAACAACTGC |
| | | Reverse | TCGGAACAGACCATCTTGC |
| Major sperm protein | A0A0V1BX53 | Forward | GCTGCTGAACCTGAAGTGAA |
| | | Reverse | ACCTCCGACCTGCTTTTCT |
| Cysteine-rich venom protein 1 | A0A0V1BM34 | Forward | CCTCCAGCGTGTTCATTTTG |
| | | Reverse | CGGCATTGACATTTTGCTTG |
| Histone H3 | A0A0V1B2J8 | Forward | TAGCAGCAAGGCAATGTCAA |
| | | Reverse | CGGCAAAAGTAAACGAACAGC |
| Receptor expression-enhancing protein 5 | A0A0V1B4B5 | Forward | AACGCCAACTGTAAGCAATGG |
| | | Reverse | GGAGTGAAAAGCCAAACAAAGA |
| Cuticlin-1 | A0A0V1BL36 | Forward | AAATGGCTGTCCGCAAGA |
| | | Reverse | TTCTCCGTCTGTTTTCAACG |
| Transmembrane matrix receptor MUP-4 | A0A0V1BTK0 | Forward | ATCGTGAGATGGGCGGAAG |
| | | Reverse | CGTGGGATAGAAAAGTCAGTGTAGAA |
| BPI2 domain-containing protein | A0A0V1B5A1 | Forward | AATCAGGGTGTGCTGTTGGA |
| | | Reverse | TGCTGCGTATGTTTCTGCTG |
| Metalloendopeptidase | A0A0V1C237 | Forward | CTGAAACCTTATCCACGAGCAG |
| | | Reverse | TGCGACGAAACACCCGTA |
| Uncharacterized protein T01_11650 | A0A0V1AZ10 | Forward | GACGCAGCATTTACAGACCA |
| | | Reverse | TTTTGGCAGTTTTCCATTCG |
| Laminin subunit beta-1 | A0A0V1B4G8 | Forward | CCAAGCAGGCGATTTTGTC |
| | | Reverse | GGTTTCGGTCGTTTCAGCA |
| Peroxidase | A0A0V1B4W8 | Forward | TTGGGCATACATTGATACAGC |
| | | Reverse | CTTGGCAGGAGTTGAAAACA |
| Glyceraldehyde 3-phosphate dehydrogenase (GAPDH) | AF452239.1 | Forward | TCACCGAAATGAAGCCTGAA |
| | | Reverse | CCCATAACCAACATAGGAGCA |

**Supplementary Table 6. Primer sequence of *T. spiralis* miRNAs for quantitation of RT-qPCR.**

| miRNA name | Primer sequence |
|---|---|
| Oligo dT adaptor primer | CAGTGCAGGGTCCGAGGTCAGAGCCACCTGGGCAATTTTTTTTTTTVN |
| tsp-miR-1 | AAGCGACCTGGAATGTAAAGAAGT |
| tsp-miR-9b-5p | AACACGCTCTTTGGTCATTTAGCTG |
| tsp-miR-32-3p | AACAAGTGCTGCAGTGTGTTTGA |
| Unk_tsp_miR-52 | AACAGTGTCACCGGTCCATTTTATC |
| tsp-miR-53-5p | AACACGCTACCCGTATCTTTCTTTC |
| tsp-miR-56 | AACAAGTACCCGTATCTCTTCTTGG |
| Unk_tsp_miR-57 | AACACGCTCACCGAATACTAAAGC |
| Unk_tsp_miR-58 | AACACGCTCACCGGATACTAAAAC |
| Unk_tsp_miR-60 | AACAAGCACCCGGATGCTAAAAC |
| Unk_tsp_miR-65 | AACAAGTACGACTGTGATTGCTCAA |
| tsp-miR-72-5p | AACAAGAGGCAAGATGTTGGCATAG |
| Unk_tsp_miR-72 | AACAAGTCGAATCGCCACATCG |
| tsp-miR-79-3p | AACACGCTAAAGCTGGATGACC |
| tsp-miR-82 | AACAAGTGAGATCACCGTGAAAGC |
| tsp-miR-86-5p | AACAAGTAAGTGAATGCTTTGCCAC |
| tsp-miR-87-3p | AACAAGGTGAGCAAAGTTTCAGGT |
| tsp-miR-100-5p | AACAGTGAACCCGTAGATCCGAA |
| tsp-miR-124-5p | AACAAGCGTGTTCACTGTCTGTC |
| tsp-miR-125b-5p | AACAAGTGGACGGATGCTCAGT |
| tsp-miR-228-5p | AACACGCAATGGCACTGGATG |
| tsp-miR-252b | AACAAGCTAAGTAGTAGTGCCGC |
| tsp-miR-263a-5p | AACAAGCTTGGCACTGTAAGAATTC |
| tsp-miR-279d-3p | AACAAGAGGCAAGATGTTGGCATAG |
| tsp-miR-375 | AACACGCTTTGTTCGTTTTGATCG |
| tsp-miR-2240c | AACAAGTGGACGGATGCTCAGT |
| tsp-miR-2496-3p | AACAAGTCACCGGGCACTAAAAC |
| tsp-miR6235-3p | AACACGCAACCCGTATTTTCCTC |
| tsp-miR-7944-3p | AACAAGTCACCGGGCACTAAATC |
| tsp-miR-9391-5p | AACACGCTCACCGGTTACTAAAAC |
| Unk_tsp_miR-1 | AACACGCTTGACTGTAATCGATTGG |

**Supplementary Table 6. continued**

| miRNA name | Primer sequence |
|---|---|
| Unk_tsp_miR-2 | AACACGCAAGATAAAGTGGAGCG |
| tsp-miR-5437 | AACACGCAAGATGAAGTGGAACG |
| Unk_tsp_miR-3 | AACACGCAATGATGAGGAGCTAAAC |
| tsp-miR3352 | ACGCCGTCTTCTTCTGTTCTATATC |
| tsp-miR-35a-3p | AACAAGTGGAAAAATTGAGCCATGG |
| tsp-miR-2238l-3p | AACAAGTGGACGGCGAATTAGTG |
| tsp-miR-107b | AACACGTGTGACTAGGCCCTG |
| tsp-miR-36-3p | AACACGCTCACCGGGTAATAATTC |
| tsp-miR-3084b-3p | AACAATTGCCAGTTCCCTCCTG |
| tsp-miR-4752 | AACAAGTCGATCATCCTTGAGAACC |
| tsp-miR7510a | AACAAGTGGGACGCTAAAGATCTTG |
| Unk_tsp_miR-37 | AACAATGTGCATGCCTGAGATCTTC |
| tsp-miR-2004 | AACAAGATGTTCCTGTGGTTGCG |
| tsp-miR7787-3p | AACAAGTCTTCGGTCTGTACTTGGT |
| Unk_tsp_miR-38 | AACAATTGTACCGCCTGGCAG |
| Unk_tsp_miR-39 | AACACGCGATTTATCAATTGTCGAA |

**Supplementary Table 7. Mature miRNAs sequence for miRNAs synthesis**

| miRNA name | Forward sequence | Reverse sequence |
|---|---|---|
| tsp-miR-1 | UGGAAUGUAAAGUAUGUAG | CCAUACUUCUUUGCAUCGCCAU |
| tsp-miR-5437 | AAGAUGAAGUGGAACGGUUAUU | UCACCGGAUCAUUUCAUCUUCU |
| tsp-miR-107 | UGACUAGGCCCUGUACAAUCCGGAU | CCAUGGUUUUGACGGUGUAUUCC |
| tsp-36-3p | UCACCGAAUUCACAGC | UGUGAAUUGUUUCCUCGGUCAUU |
| tsp-miR-181b-5p | UUCAGUGCUGACGGCGAACAAU | UGUUGCCUUCAAUACUGAAAGA |
| Negative control | GGUUCGUACGUACACUGUUCA | UGAACAGUGUACGUACGAACC |
